# Dendritic cells type 1 control the formation, maintenance, and function of tertiary lymphoid structures in cancer

**DOI:** 10.1101/2024.12.27.628014

**Authors:** Raphaël Mattiuz, Jesse Boumelha, Pauline Hamon, Jessica Le Berichel, Abishek Vaidya, Brian Y. Soong, Laszlo Halasz, Emir Radkevich, Hye Mi Kim, Matthew D. Park, Romain Donne, Leanna Troncoso, Darwin D’Souza, Medard Ernest Kaiza, Ian P. MacFawn, Meriem Belabed, Guillaume Mestrallet, Etienne Humblin, Raphaël Merand, Clotilde Hennequin, Giorgio Ioannou, Sinem Ozbey, Igor Figueiredo, Samarth Hegde, Alexander Tepper, Hajer Merarda, Erika Nemeth, Simon Goldstein, Amanda M. Reid, Moataz Noureddine, Alexandra Tabachnikova, Jalal Ahmed, Alexandros D. Polydorides, Nina Bhardwaj, Amaia Lujambio, Zhihong Chen, Edgar Gonzalez Kozlova, Seunghee Kim-Schulze, Joshua D. Brody, Michael Schotsaert, Christine Moussion, Sacha Gnjatic, Catherine Sautès-Fridman, Wolf Herman Fridman, Vladimir Roudko, Brian D. Brown, Thomas U. Marron, Jason G. Cyster, Hélène Salmon, Tullia C. Bruno, Nikhil S. Joshi, Alice O. Kamphorst, Miriam Merad

## Abstract

Tertiary lymphoid structures (TLS) are organized immune cell aggregates that arise in chronic inflammatory conditions. In cancer, TLS are associated with better prognosis and enhanced response to immunotherapy, making these structures attractive therapeutic targets. However, the mechanisms regulating TLS formation and maintenance in cancer are incompletely understood. Using spatial transcriptomics and multiplex imaging across various human tumors, we found an enrichment of mature dendritic cells (DC) expressing high levels of CCR7 in TLS, prompting us to investigate the role of DC in the formation and maintenance of TLS in solid tumors. To address this, we developed a novel murine model of non-small cell lung cancer (NSCLC) that forms mature TLS, containing B cell follicles with germinal centers and T cell zones with T follicular helper cells (T_FH_) and TCF1^+^PD-1^+^ progenitor exhausted CD8^+^ T cells (Tpex). Here we show that, during the early stages of tumor development, TLS formation relies on IFNγ-driven maturation of the conventional DC type 1 (cDC1) subset, their migration to tumor-draining lymph nodes (tdLN), and recruitment of activated T cells to the tumor site. As tumors progress, TLS maintenance becomes independent of T cell egress from tdLN, coinciding with a significant reduction of cDC1 migration to tdLN. Instead, mature cDC1 accumulate within intratumoral CCR7 ligand-enriched stromal hubs. Notably, timed depletion of cDC1 or disruption of their migration to these stromal hubs after TLS are formed alters TLS maintenance. Importantly, we found that cDC1-mediated antigen presentation to both CD4^+^ and CD8^+^ T cells and intact CD40 signaling, is critical for the maintenance of TLS, the preservation of the T_FH_ cell pool, the formation of germinal center and the production of tumor-specific IgG antibodies. These findings underscore the key role of mature cDC1 in establishing and maintaining functional TLS within tumor lesions and highlight the potential for cDC1-targeting therapies as a promising strategy to enhance TLS function and improve anti-tumor immunity in patients with cancer.

## Main

Tertiary lymphoid structures (TLS) are organized immune aggregates that resemble canonical secondary lymphoid organs, containing naïve and antigen-experienced T and B cells organized in T cell zones and B cell follicles, respectively (Fridman et al., 2023; Teillaud et al., 2024). With few exceptions, patients with tumors enriched in TLS have improved outcome and enhanced response to immune checkpoint blockade (ICB) (Cabrita et al., 2020; Dieu-Nosjean et al., 2008; Goc et al., 2014; Helmink et al., 2020; Petitprez et al., 2020), suggesting that TLS may contribute to the priming of naïve T cells or enable the reactivation of effective T cell responses upon ICB treatment (Schumacher and Thommen, 2022).

Previous findings have reported that TLS in non-small cell lung cancer (NSCLC) are associated with DC-LAMP^+^ mature dendritic cells (DC) (Goc et al., 2014). However, the exact role for antigen presentation in the maintenance and functionality of TLS remain unclear (Schumacher and Thommen, 2022). Conventional dendritic cells (cDC) are uniquely equipped to prime and educate T cells (Durai and Murphy, 2016). Two subsets of cDC have been identified in human and mice including cDC1, which excel in the cross-presentation of cell-associated antigens to CD8^+^ and CD4^+^ T cells and cDC2 that are most potent to present soluble proteins to CD4^+^ T cells (Durai and Murphy, 2016; Guilliams et al., 2014). Using spatial transcriptomics and multiplex imaging analysis of various human tumors, we found that pathology-annotated TLS are composed of cDC in a mature molecular state along with T follicular helper cells (T_FH_), TCF1^+^PD-1^+^ progenitor exhausted CD8^+^ T cells (Tpex), and B cells. Together, these results prompted us to explore the contribution of cDC to the formation and maintenance of TLS in tumor lesions.

Given that cDC can both prime T cells within tumor-draining lymph nodes (tdLN) and reactivate T cells at the tumor site, it became essential to develop models that allow for the conditional manipulation of cDC to dissect their specific contributions in these distinct tissue sites. To address this, we engineered a tumor model that reliably forms mature TLS and developed strategies for constitutive or conditional depletion or genetic manipulation of cDC at different timepoints following tumor development. Here, we show that mature DC, particularly mature cDC1, are central to TLS formation, maintenance, and function in tumors.

### Mature DC are enriched in human TLS, where they interact with T_FH_ and B cells across several tumor types

Mature DC exhibit a discrete cellular program that is upregulated in cDC1 and cDC2 upon capture of cell debris in tumor lesions and reflect the state in which cDC interact with T cells (Belabed et al., 2023; Maier et al., 2020). To map the spatial distribution of cDC subsets and states within human tumors, we performed a spatial transcriptomics (Visium 10x Genomics) analysis of human non-small cell lung cancer (NSCLC), hepatocellular carcinoma (HCC), colorectal cancer (CRC), and clear cell renal cell cancer (ccRCC) (Meylan et al., 2022). We found a significant enrichment of mature DC (*CCR7, FSCN1, CD40, CD80, CCL17, CCL21*), cDC1 (*XCR1, CLNK, CADM1*), T_FH_ (*CXCL13*, *TCF7, ICOS, CD40LG, PDCD1*), naïve T cells (*IL7R, CCR7, TCF7*), memory B cells (*CD19, BANK1*) and germinal center B cells (*LMO2, MEF2B, RGS13*) molecular programs were enriched in pathology-annotated TLS but were markedly depleted at the tumor core (**Fig. 1a, b and Extended data Fig. 1a-c**). In contrast, monocytes and macrophages were distributed both within and outside TLS (**Extended data Fig. 1b**). We also examined gene expression pathways within these mature DC – either by characterizing those found in TLS spots from spatially-mapped human tumor lesions (**Extended data Fig. 1d**) or those identified in TLS-enriched NSCLC and HCC lesions by scRNA-seq (Leader et al., 2021; Magen et al., 2023) (**Supplementary Table 1**). In both analyses, we found that mature DC in TLS-enriched lesions upregulated MHC-II and MHC-I antigen presentation machinery and CD40 signaling pathways **(Fig. 1c)**, suggesting interactions with T cells.

**Fig 1.**
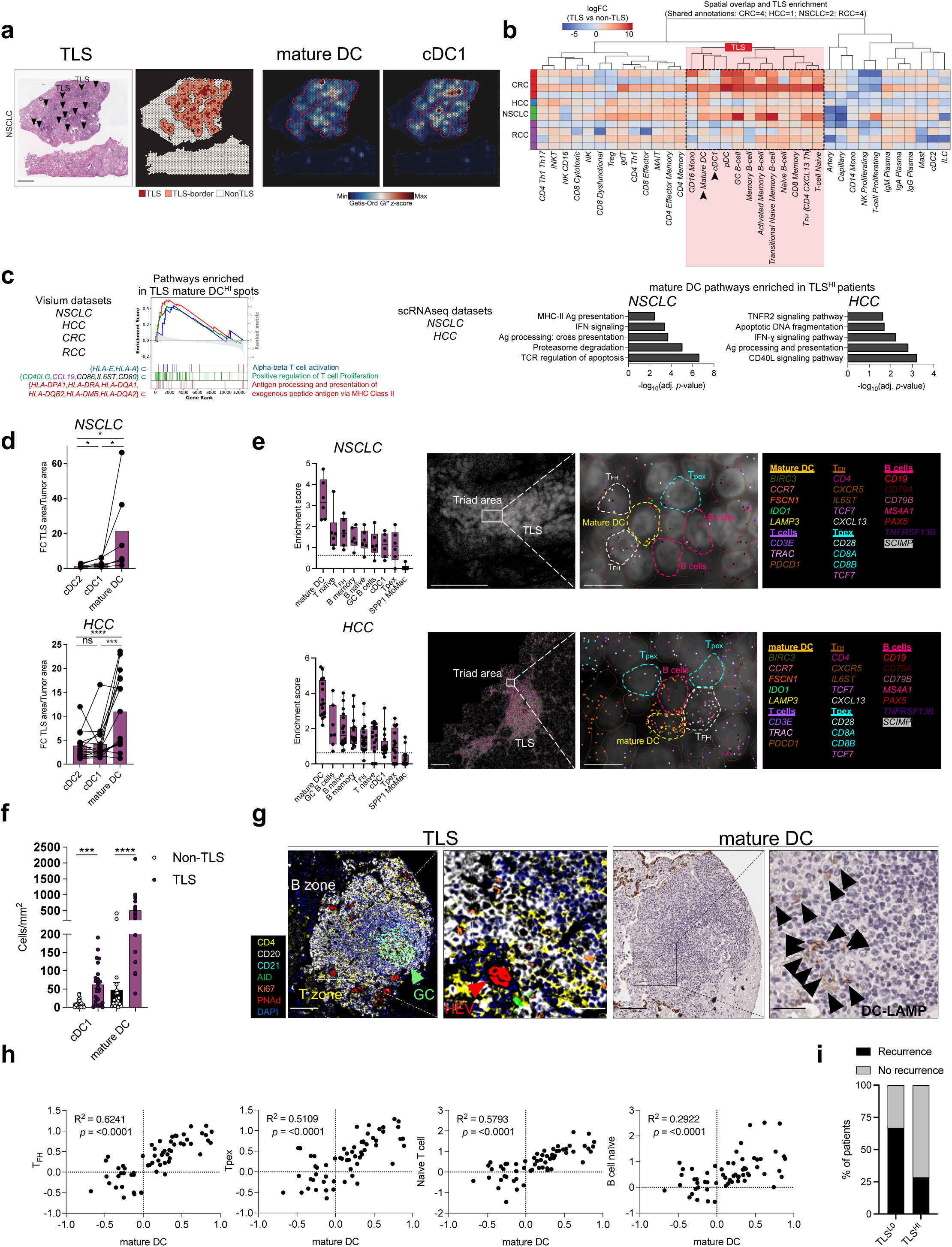
Mature DC accumulate in TLS across human cancers. **a, b** Formalin-fixed paraffin-embedded (FFPE) Visium Spatial Transcriptomics analysis of NSCLC (*n* = 2), HCC (*n* = 1), CRC (*n* = 4), and RCC tumors (*n* = 4, data from *Meylan et al., Immunity, 2022*). **a** Representative Visium NSCLC sample showing pathology-annotated TLS (left, scale bar, 1mm), TLS and border spots (two layers of spots located outside the TLS), and hotspots for mature DC and cDC1 gene signatures (right). **b** Heatmap displaying log fold change (logFC) in TLS spots versus non-TLS spots across shared immune cell types, deconvoluted from scRNA-seq references and segmented across tumor types. **c** Gene set enrichment analyses of TLS mature DC-high Visium spots compared to TLS mature DC-low spots across tumor types (left) and of mature DC scRNA-seq clusters from TLS-high patients compared to others in NSCLC and HCC (right) (*n* = 14 TLS^HI^ and *n* = 18 TLS^LO^ NSCLC patients; *n* = 8 TLS^HI^ and *n* = 6 TLSL^LO^ HCC patients). **d, e** MERFISH Spatial Transcriptomics analysis of NSCLC (*n* = 6) and HCC (*n* = 16) tumors. **d** Fold change of mature DC, cDC1, and cDC2 populations in TLS versus non-TLS areas in NSCLC (top) and HCC (bottom). \**p* < 0.05; \*\*\**p* < 0.001; \*\*\*\**p* < 0.0001 (Wilcoxon rank sum test). **e** Cell proximity analysis showing enrichment score for immune populations within 30 µm of mature DC in TLS areas, with representative TLS area illustrating interactions between mature DC, T_FH_, Tpex, and B cells in NSCLC (top, scale bars, left 100µm and right 10µm) and HCC (bottom, scale bars, left 200µm and right 10µm). **f** Quantification of mature DC (DC-LAMP^+^) and cDC1 (CLEC9A^+^) densities in non-TLS versus TLS regions by multiplex IHC (MICSSS) in NSCLC patients (*n* = 23); epithelial cells expressing DC-LAMP were excluded from the analysis. \*\*\**p* < 0.001; \*\*\*\**p* < 0.0001 (paired *t*-test). **g** Representative multiplex IF staining of a mature TLS (CD4, CD20, CD21, AID, Ki67, PNAd, DAPI), paired with consecutive DC-LAMP IHC staining of NSCLC tumors. Representative of 14 TLS. Scale bars, 100µm (overview) and 50µm (zoom). **h** Gene signatures derived from bulk transcriptomes paired with TLS annotations in NSCLC, showing correlations of bulk RNA-seq gene signatures between mature DC and T_FH_, Tpex, naïve T and B cells in TLS-positive patients (*n* = 60, from Genentech POPLAR dataset). Gene signatures for different immune cell types were generated from prior literature (*Magen et al., 2023*). *p* < 0.0001 (Pearson correlation). **i** Recurrence status in TLS-high versus TLS-low groups in a neoadjuvant anti-PD-1 trial for patients with HCC (*n* = 13 patients).

To investigate mature DC cellular interactions within TLS at the single-cell level in NSCLC and HCC, we employed a targeted spatial transcriptomic technology (MERFISH) to map 500 gene probes that we had previously identified to be highly enriched in tumor lesions by scRNA-seq (Grout et al., 2022; Leader et al., 2021; Magen et al., 2023) (**Supplementary Table 2**). We confirmed that mature DC were significantly enriched in TLS within the tumor, compared to their immature counterparts (cDC1 and cDC2) **(Fig. 1d).** Proximity analysis revealed that within TLS, mature DC were closely located (within a 30 μm radius) to naïve B and T cells, germinal center B cells, and T_FH_ and Tpex cells **(Fig. 1e and Extended data Fig. 1e-g)**. DC-LAMP, encoded by *LAMP3*, was found to be exclusively expressed by mature DC within the immune compartment of NSCLC and HCC by scRNA-seq **(Extended data Fig. 1h)**. Using multiplex imaging, we confirmed at the protein level that whilst both DC-LAMP^+^ mature DC and CLEC9A^+^ cDC1 were enriched in TLS, mature DC were more abundant **(Fig. 1f, Extended data. Fig.1i, Supplementary Table 3**). We also found that mature DC were primarily located in the T cell zone or at the T-B zone border close to high endothelial venules (HEV) similar to their positioning in lymph nodes (Moussion and Girard, 2011) **(Fig. 1g, Extended Fig. 1j)**. Importantly, we observed a similar density of mature DC to TLS with or without germinal centers (**Extended data Fig. 1k**).

In patients with locally advanced or metastatic NSCLC who failed platinum therapy, a randomized phase II trial (POPLAR) comparing PD-L1 blockade (atezolizumab) with chemotherapy (Docetaxel) reported a significant survival benefit of PD-L1 blockade (atezolizumab) over chemotherapy (Docetaxel) (Patil et al., 2022). Bulk RNA-seq analysis of pre-treatment biopsies within this cohort showed that those with higher TLS density, as annotated by a pathologist, exhibited a robust mature DC gene signature and superior survival outcomes (**Extended data Fig. 1l, m**). Furthermore, in these patients, the mature DC signature correlated with signatures of components of TLS including T_FH_, Tpex, and naïve B and T cells (**Fig. 1h and Extended data Fig. 1n, Supplementary Table 4**). These observations suggest that enrichment in mature DC correlates with TLS occurrence and can serve as a prognostic marker for improved survival. Furthermore, in a cohort of HCC patients treated with neo-adjuvant PD-1 blockade (Marron et al., 2022), post-treatment tumor resections enriched in TLS also had increased infiltration of mature DC and lower recurrence rates **(Fig. 1i and Extended data Fig. 1o)**. Collectively, these findings underscore the potential key role of mature DC within TLS in fostering the response to ICB in human tumors, prompting our interest in understanding the contribution of mature DC in TLS formation, maintenance and function in tumors.

### Mature cDC1 accumulate within TLS in preclinical lung cancer tumors

To study the role of cDC in the formation and function of TLS in cancer, we used a modified murine lung adenocarcinoma (LUAD) line, named KP-HELLO-2, derived from *KRAS^LSL-G12D^*^;^*Trp53^fl/fl^*(KP) mice (Damo et al., 2023) expressing the model antigen HELLO (fusion neoantigen of the B cell antigen HEL, the CD8 antigen GP33, and the CD4 antigen GP66) (Cui et al., 2021) and the Thy-1.1 antigen (CD90.1) **(Fig. 2a)**. Intravenous injection of KP-HELLO-2 cells led to the formation of orthotopic lung adenocarcinoma lesions with high numbers of TLS **(Fig. 2b and Extended data Fig. 2a),** many of which displayed prominent germinal centers that harbor CD21^+^ stromal follicular dendritic cells adjacent to activation-induced cytidine deaminase (AID)^+^ germinal center B cells (**Fig. 2c**). Consistent with what we observed in human tumor lesions, murine TLS were enriched in T_FH_ (CD8^-^TCF1^+^PD-1^+^), Tpex (CD8^+^TCF1^+^PD-1^+^) (**Fig.2d**), and tumor antigen-specific CD4^+^ T cells (CD4^+^CD45.1-SMARTA^+^) and B cells (B220^+^HEL^+^) (**Fig. 2e**). Furthermore, DC expressing high levels of CCR7, MHC-II, and CD86 accumulated in TLS, where they interacted with CD4^+^ T cells, CD8^+^ T cells and B cells (**Fig. 2f**).

**Fig 2.**
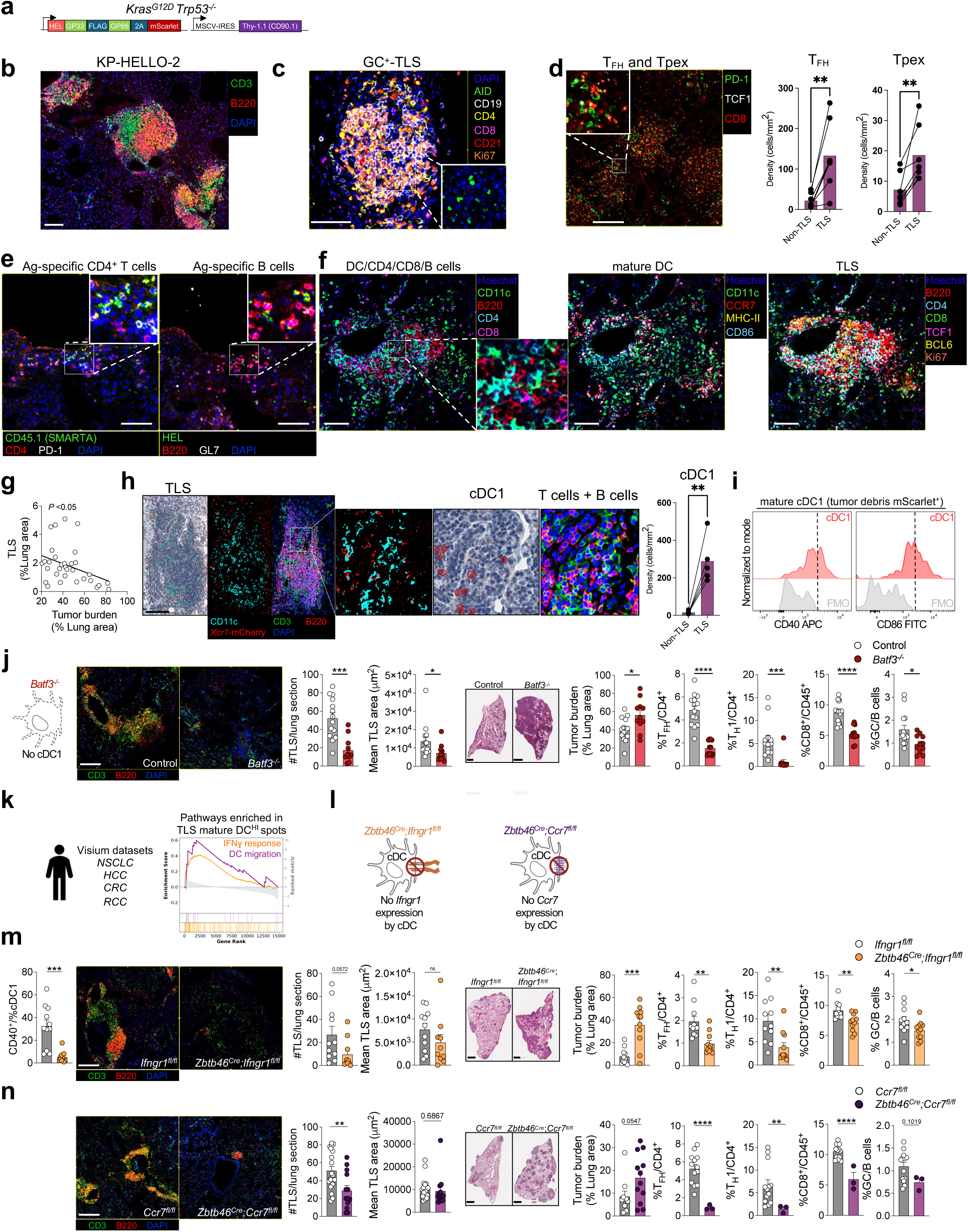
Mature cDC1 control TLS formation. **a** Schematic representation of constructs used to generate KP-HELLO-2. **b-h** Immunofluorescence analysis of representative KP-HELLO-2 TLS at d15 post-engraftment in WT mice. **b** Staining of B cell and T cell zones with B220 and CD3. Scale bar, 100µm. **c** GC^+^-TLS stained with AID, CD19, CD4, CD8, CD21, and Ki67. Representative of 7 mice. Scale bar, 50µm. **d** Representative images and density quantification of T_FH_ (CD8^-^ PD-1^+^ TCF1^+^) and Tpex (CD8^+^ PD-1^+^ TCF1^+^) in TLS and non-TLS areas. Scale bar, 200µm. \*\**p* < 0.01 (paired *t*-test). **e** Adoptively transferred antigen-specific CD4 T cells (SMARTA CD45.1) stained with CD45.1, PD-1, and CD4; antigen-specific B cells stained with HEL antigen, B220, and GL7. Representative of 10 mice. Scale bar, 100µm. **f** mature DC (CD11c^+^ MHC-II^+^ CCR7^+^ CD86^+^) within mature TLS stained with B220, CD4, CD8, TCF1, BCL6, and Ki67. Representative of 2 mice. Scale bar, 100µm. **g** Inverse correlation between TLS area and tumor area (% of lung area) in WT mice (*n* = 32). **h** Representative images of TLS from *Xcr1-mCherry-Cre* mice showing CD3 and B220 immunofluorescence, paired with consecutive IHC staining for CD11c and mCherry, and quantification of cDC1 density in TLS and non-TLS areas (*n* = 5 mice). Scale bar, 100µm. **i** cDC1 maturation status (CD40 and CD86 expression in mScarlet^+^ cDC1) at d12 post-engraftment (representative of 13 mice from two experiments). Immunofluorescence of TLS (CD3, B220, DAPI, scale bar, 200µm), number and mean size, tumor area assessed by H&E on consecutive sections, scale bar, 1mm, and intratumor quantification of T(PD-1^+^ CXCR5^+^ CD4^+^), GC B cells (CD19^+^ IgD^-^, IgM^-^, GL7^+^), T_H_1 (IFNγ^+^ CD4^+^), and CD8^+^T cells by flow cytometry on d15 in **j** *Batf3^-/-^* (*n* = 11-14 per group, pooled from two independent experiments) and **m** *Zbtb46^Cre^; Ifngr1^fl/fl^* (*n* = 10-11 per group, representative of two independent experiments) and **n** *Zbtb46^Cre^; Ccr7^fl/fl^* mice (*n* = 13-20 per group, pooled from two independent experiments) versus littermate controls. **k** Gene set enrichment analysis for IFNγ response and DC migration in TLS mature DC-high Visium spots compared to other spots in human NSCLC, HCC, CRC, and RCC slides. **l** Schematic representation of *Ifngr1* and *Ccr7* deletion in DC. \**p* < 0.05; \*\**p* < 0.01; \*\*\**p* < 0.001; \*\*\*\**p* < 0.0001 (unpaired *t*-test).

TLS are inversely correlated with tumor burden, suggesting that these structures contributed to anti-tumor immunity **(Fig. 2g)**. Imaging analysis of tumor lesions in cDC1-reporter mice confirmed that cDC1 (CD11c^+^*Xcr1^mCherry+^*) accumulated in TLS in proximity to T and B cells **(Fig. 2h),** as observed in human cancer lesions and most cDC1 were in a mature state (CD40^+^CD86^+^) and charged with tumor antigens (mScarlet^+^) (**Fig. 2i).**

To probe whether cDC or specific cDC subsets played a role in TLS formation, we injected KP-HELLO-2 cells into *Batf3^-/-^* mice that specifically lacked cDC1 but not cDC2 **(Extended data Fig. 2b).** Absence of cDC1 led to significantly reduced TLS in both numbers and size, and reduced T_FH_ (CXCR5^+^PD-1^+^ CD4^+^ T cells), T_H_1 (IFNγ^+^ CD4^+^ T cells), CD8^+^ T cells, and germinal center B cells within tumors which was accompanied with increased tumor burden **(Fig. 2j, Supplementary Table 5)**. cDC1 were also required for TLS formation in mice challenged with orthotopic Hep-53.4 liver tumors and orthotopic AKPS colon tumors **(Extended data Fig. 2c)**. Notably, mature cDC1 generated *in vitro* upon capture of apoptotic KP-HELLO-2 tumor cells were also sufficient to drive the formation of immune aggregates *ex vivo* upon co-culture with purified intratumoral T and B cells within tumor spheroids **(Extended data Fig. 2d)**.

### CCR7 and IFNγ receptor expression on cDC is critical for TLS formation

We next sought to investigate the cDC molecular signals that regulate TLS formation in tumors. Using the Visium spatial transcriptomics platform to quantify gene signature scores in mature DC-enriched TLS spots, we observed an enrichment in IFNγ gene signature **(Fig. 2k).** IFNγ receptor (IFNγR) signaling promotes DC immunogenic function and their ability to instruct effective anti-tumor T cell responses (Garris et al., 2018; Maier et al., 2020; Mattiuz et al., 2021). To examine whether IFNγ signaling in DC plays a role in TLS formation, we generated mice lacking IFNγR specifically within the DC compartment (*Zbtb46^Cre^;Ifngr1^fl/fl^* mice) (**Fig. 2l**). We found that absence of IFNγR signaling prevented DC maturation (CD40^+^), reduced TLS formation and increased tumor burden, which was accompanied by a marked decrease in T_FH_, T_H_1, CD8^+^ T cells, and germinal center B cells within tumors **(Fig. 2m)**.

CCR7 upregulation is required for DC migration to the T zone of secondary lymphoid organs towards CCR7 ligands produced by the lymphatic endothelium and stromal cells that populate the T zone (Luther et al., 2000; Ohl et al., 2004), which helps position DC charged with tissue antigens in proximity of naïve T cells, thus enabling the priming of tumor-specific immunity. To probe whether DC migration to T zones of lymphoid organs contributed to TLS formation, we generated mice lacking *Ccr7* specifically within the DC compartment (*Zbtb46^Cre^;Ccr7^fl/fl^* mice) (**Fig. 2l**) and challenged these mice with KP-HELLO-2 tumors. Importantly, CCR7 deletion in DC significantly disrupted TLS development and increased tumor burden with a notable decrease in T_FH_, T_H_1, CD8^+^ T cells, and germinal center B cells in tumors **(Fig. 2n)**. Altogether, these results establish that IFNγ-driven maturation and DC migration to lymphoid structures contribute to the formation of TLS in lung cancer lesions.

### cDC1 are required locally for TLS maintenance

cDC are required for T cell priming in tdLN and efficient priming of T cells may contribute to the formation of TLS, but cDC also interact with T cells within tumor tissues and may contribute to TLS maintenance locally. To examine whether TLS formation is dependent on cDC within tdLN or at the tumor site, we first explored the contribution of T cells primed in tdLN to TLS formation using FTY720 to inhibit lymphoid egress from the tdLN (Matloubian et al., 2004). We found that inhibition of T cell egress during the first 9 days following tumor engraftment significantly reduced TLS formation, whereas inhibition of T cell egress after day 9 had little impact on the number of TLS in tumors **(Fig. 3a)**. These results establish that T cell priming in the tdLN is critical for TLS formation but not for TLS maintenance locally in tumor lesions, prompting us to measure longitudinal changes in immune cell composition that were associated with TLS formation and maintenance in tumors.

**Fig 3.**
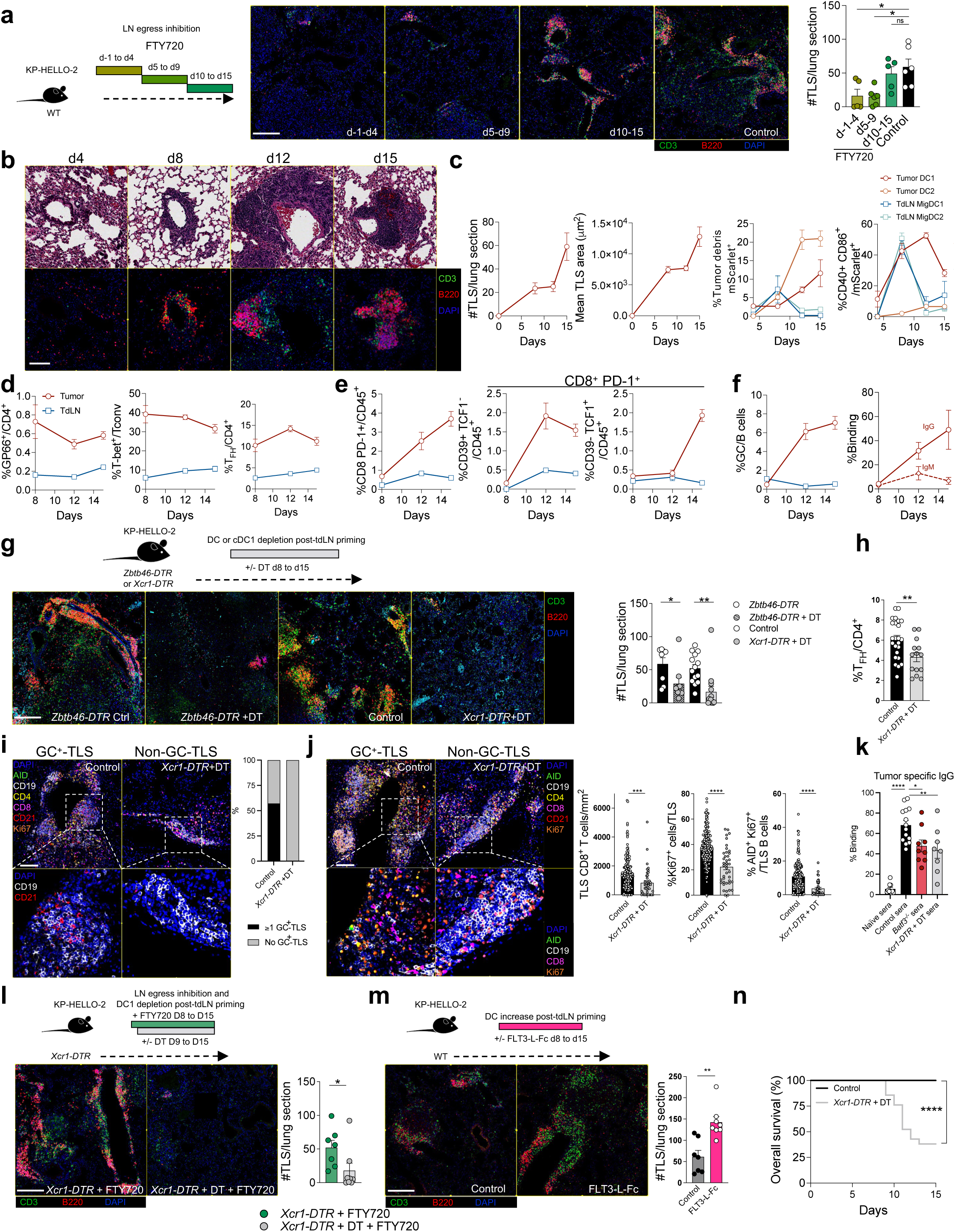
Mature cDC1 are required locally for TLS maintenance and function. **a-c, g, i, j, l, m** Immunofluorescence analysis of TLS (CD3, B220, DAPI staining) and quantification at d15 or as indicated. **a** T cell egress from tdLN blocked with FTY720 from d-1 to d4, d5 to d9, or d10 to d15 post-tumor engraftment. TLS quantification was performed at day 15 (*n* = 5-6 per group, representative of two independent experiments). Scale bar, 200µm. **b, c** TLS formation kinetics (*n* = 6-7 per timepoint, representative of two independent experiments), with **b** H&E staining on consecutive sections showing vessels, scale bar, 100µm, and **c** kinetic analysis of immune populations in tumor and tdLN by flow cytometry, including percentages of DC uptaking tumor debris (mScarlet^+^) (left) and their maturation (CD40^+^ CD86^+^) (right). **d** Tumor-antigen-specific, T-bet^+^, and T_FH_ (PD-1^+^ CXCR5^+^) cells among CD4^+^ T cells. **e** CD8^+^ PD-1^+^ T cells, subdivided into CD39^+^ TCF1^-^ and CD39^-^ TCF1^+^ populations, among immune cells. **f** GC B cells among B cells (left) and kinetics of serum IgG and IgM binding to tumor cells *ex vivo* at the indicated time points (right). **g-k** DC or cDC1 depletion using DT from d8 to d15 in DTR mice. **g** TLS quantification in *Zbtb46-DTR*, *Xcr1-DTR* and control mice treated or not with DT (*n* = 8-14 per group, pooled from two independent experiments). Scale bar, 200µm. **h** Quantification of intratumor T_FH_ cells among CD4^+^ T cells at d15 in *Xcr1-DTR* mice treated with DT versus control mice (*n* = 14-25 per group, pooled from four independent experiments). **i, j** GC^+^-TLS staining (AID, CD19, CD4, CD8, CD21, and Ki67) in *Xcr1-DTR* mice treated with DT versus control mice, **i** with zoom depicting follicular DC (CD19^-^ CD21^+^) within GC^+^-TLS and proportions of mice with ≥1 GC^+^-TLS (*n* = 3-7 per group, 196 TLS analyzed, including 7 GC^+^-TLS) per whole lung slide. Scale bar, 100µm. **j** AID, Ki67 and CD8 staining within TLS and quantification of CD8^+^ T cells, Ki67^+^ cells, and AID^+^ Ki67^+^ B cells in TLS of *Xcr1-DTR* mice treated with DT versus control mice (*n* = 3 per group, 229 TLS analyzed). Scale bar, 100µm. **k** Serum IgG binding to tumor cells *ex vivo* from naïve, control, *Batf3^-/-^*, and *Xcr1-DTR* treated with DT mice (*n* = 8-14 per group, pooled from two independent experiments). Sera were collected at d15. **l** TLS quantification in *Xcr1-DTR* mice treated with FTY720 starting at d8 and with or without DT starting at d9 (*n* = 6-8 per group). Scale bar, 200µm. **m** TLS quantification in WT mice treated or not with FLT3-L-Fc at d8 (*n* = 7-8 per group, representative of two independent experiments). Scale bar, 200µm. **a, g, h, j-m** \**p* < 0.05; \*\**p* < 0.01; \*\*\**p* < 0.001; \*\*\*\**p* < 0.0001 (unpaired *t*-test). **n** Overall survival of *Xcr1-DTR* mice treated with DT (from d8 to d15) versus control mice (*n* = 21-29 per group, pooled from three independent experiments), with all mice reaching the humane endpoint by d15 post-engraftment. \*\*\*\**p* < 0.0001 (Log-rank (Mantel-Cox) test).

Longitudinal profiling of tumor lesions revealed that immature TLS arise from or near the vasculature and progress towards mature TLS with the formation of B cell follicles and T cell zones at later time points (**Fig. 3b**). Specifically, we observed a drastic increase in TLS number and size during tumor progression and the increase in TLS was particularly steep after day 8 **(Fig. 3c)**. Interestingly, the number of mature cDC1 and cDC2 carrying tumor-associated antigens (mScarlet^+^) increased in the tdLN until day 8 following tumor implantation, after which their accumulation in the tdLN was strongly reduced while instead they started to accumulate locally at the tumor site (**Fig. 3c**). Tumor-specific CD4^+^, T_FH_, and T_H_1 populations were maintained over time, while there was a steady accumulation of antigen-experienced (PD-1^+^) and terminally exhausted (CD39^+^TCF1^−^PD-1^+^) CD8^+^ T cells during tumor progression (**Fig.3e**). At later timepoints, coinciding with the greatest expansion of TLS, we also observed a striking expansion of Tpex (CD39^−^TCF1^+^PD-1^+^), accompanied by a significant increase in the frequency of germinal center B cells and class-switched tumor-specific IgG antibodies (**Fig. 3f**). Altogether, these observations reveal that the accumulation of mature cDC1 and cDC2 in tumors correlate with the accumulation of Tpex, germinal center B cells, IgG antibody production and TLS expansion at the tumor site, prompting us to explore the role of tumor infiltrating cDC in TLS maintenance locally.

To probe the role of cDC in TLS maintenance, we generated two models to either deplete the entire cDC compartment or specifically cDC1 starting at day 8 post tumor implantation to allow for T cell priming, egress from the tdLN, and accumulation in tumor tissues. We reconstituted wildtype mice with bone marrow cells isolated from mice expressing the diphtheria toxin receptor (DTR) within ZBTB46^+^ cDC (*Zbtb46-DTR*), which enables restricted cDC depletion after DT injection while sparing endothelial cells, which also express *Zbtb46* (Loschko et al., 2016) **(Extended data Fig. 3a)**. We also generated *Xcr1-DTR* mice by crossing *Xcr1^Cre^* mice with *Rosa26^lox−stop−lox-hDTR^* mice to enable timed deletion of XCR1^+^ cDC1 while sparing cDC2 (Ferris et al., 2020; Mattiuz et al., 2021, 2018) **(Extended data Fig. 3a)**. Strikingly, we found that the depletion of the entire cDC compartment, or specifically the cDC1 subset, at day 8 following tumor engraftment significantly reduced TLS numbers in tumor tissues at similar levels **(Fig. 3g),** prompting us to focus mainly on the role of cDC1 in TLS maintenance in tumor-bearing mice.

Importantly, cDC1 depletion reduced the frequency of T_FH_ cells in tumors (**Fig. 3h)**, led to a disappearance of germinal centers (**Fig. 3i**), reduced the number of CD8^+^ T cells and proliferating cells within TLS, and reduced the number of AID^+^ Ki67^+^ germinal center B cells (**Fig. 3j)** and tumor-specific IgM and class-switched IgG antibodies **(Fig. 3k and Extended data Fig. 3b)**. Altogether, these results established that cDC1 are required locally for the maintenance and function of TLS.

To further confirm that cDC1 promote TLS maintenance through local interaction with tumor infiltrating T cells, we blocked late-stage tdLN egress after priming occurred. We found that while blocking T cell egress at day 8 following tumor implantation did not affect TLS number, concomitant cDC1 depletion significantly reduced TLS maintenance in tumors **(Fig. 3l)**. Accordingly, we found that administration of the DC growth factor FMS-like tyrosine kinase 3 ligand agonist fused to an Fc domain (FLT3L-Fc) on day 8 post tumor implantation significantly increased the number of cDC and cDC1 (**Extended data Fig. 3c**) and doubled TLS numbers in tumors **(Fig. 3m)**. Collectively, these results indicate that TLS maintenance and function in tumor tissues depended on tumor-infiltrating mature cDC1. In line with the role of TLS in anti-tumor immunity and the requirement of cDC1 for TLS maintenance, depletion of cDC1 at day 8 post-tumor implantation significantly reduced overall survival of tumor-bearing mice (**Fig. 3n**).

### cDC1 migration to CCR7 ligands-enriched stromal hubs and sustained antigen presentation locally control TLS maintenance

The observation that cDC significantly altered their migration trajectory to accumulate in tumor tissues rather than in the tdLN after day 8 post tumor-engraftment suggested that they were actively recruited to a tissue hub. To explore the molecular signals that promoted cDC migration to specific tumor sites and their contribution to TLS maintenance locally, we generated a ligand-receptor analysis map across the different cellular compartments that populate the TLS in human NSCLC lesions using MERFISH probes (**Fig. 4a**). We found that mature DC within TLS expressed the highest levels of ligand-receptor pairs that interact with CD4^+^ T cells, including PD-L1 (*CD274*), PD-L2 (*PDCD1LG2*), *ICOSLG,* CD80/86, CCR4 ligands (*CCL17, CCL22*) and CD40.

**Fig 4.**
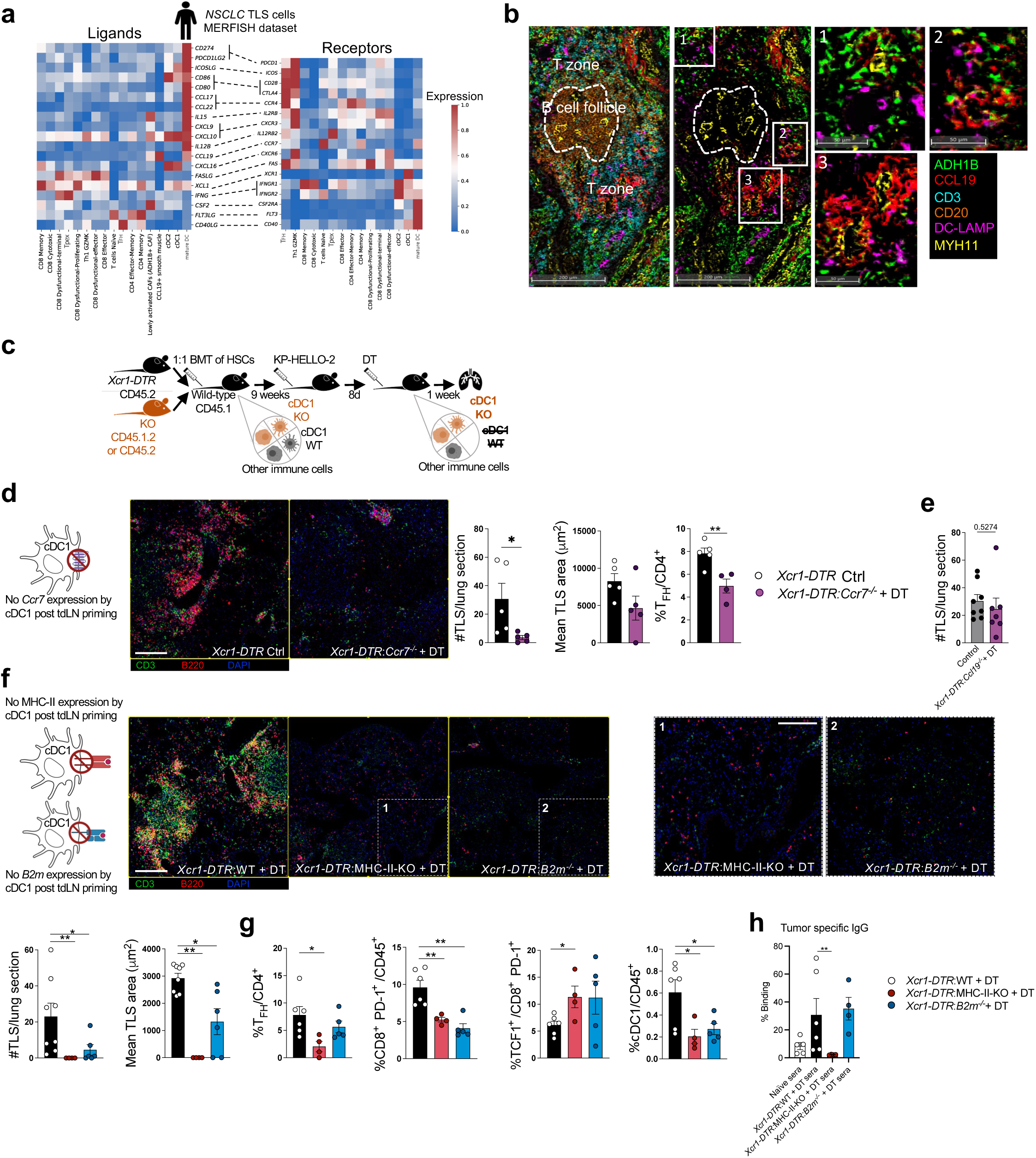
Local cDC1 Migration to CCR7 Ligand Hubs and Antigen Presentation control TLS Maintenance. **a** Heatmap showing relative expression of ligand-receptor pairs by MERFISH between ADH1B^+^ CAF, CCL19 perivascular cells, DC and T cell populations within TLS in NSCLC patients (*n* = 6). **B** Representative multiplex IHC staining of a TLS from an NSCLC tumor (stained for ADH1B, CCL19, CD3, CD20, DC-LAMP, and MYH11), representative of 3 patients. Panel 1 shows CCL19 with both ADH1B^+^ and MYH11^+^ cells, panel 2 shows CCL19 co-localizing with ADH1B^+^ cells and panel 3 shows CCL19 around MYH11^+^ cells. Scale bars, 200µm (overview) and 50µm (zoom). **C.** Schematic of *Xcr1-DTR*: KO 1:1 mixed bone marrow chimeras experiments, DT administration starting at d8 post-tumor engraftment resulted in the depletion of the entire wild-type cDC1 compartment, leaving only cDC1 that are knockout for the genes of interest. **d-g** Immunofluorescence analysis of TLS (stained for CD3, B220, and DAPI), with quantification of TLS number and mean area at d15. **D** Comparison of *Xcr1-DTR* controls versus DT-treated *Xcr1- DTR:Ccr7^-/-^* CD45.1.2 mice (*n* = 5 per group, representative of two independent experiments). Scale bar, 200µm. **e** Comparison of control bone marrow chimera versus DT-treated *Xcr1-DTR:Ccl19^-/-^* mice (*n* = 7-8 per group). **F** Comparison of DT-treated *Xcr1-DTR*/WT versus *Xcr1-DTR*:MHC-II-KO and *Xcr1-DTR:B2m^-/-^*mice (*n* = 4-8 per group, representative of two independent experiments). Panel 1 and 2 depicting B cell and T cell dispersion (right). Scale bars, 200µm (overview) and 100µm (zoom). Intratumor quantification by flow cytometry at d15 includes **d**, **g** T_FH_ among CD4^+^ T cells, **g** CD8^+^ PD-1^+^ T cells among immune cells, TCF1^+^ cells among CD8^+^ PD-1^+^ T cells, and cDC1 among immune cells. **H** Serum IgG binding to tumor cells *ex vivo* from naïve, *Xcr1-DTR*:WT, *Xcr1-DTR*:MHC-II-KO and *Xcr1- DTR:B2m^-/-^* treated with DT mice (*n* =4-6 per group, representative of two independent experiments). Sera were collected at d15. **D, e, g** \**p* < 0.05; \*\**p* < 0.01 (unpaired *t*-test). **f, h** \**p* < 0.05; \*\**p* < 0.01 (Mann-Whitney U test).

Mature DC significantly upregulated *CCR7*, while we have previously shown that one of CCR7 ligands, namely *CCL19,* was highly expressed by ADH1B^+^ cancer-associated fibroblasts (CAFs) and perivascular cells (Grout et al., 2022). Accordingly, we found that in human NSCLC, TLS formed around CCL19-producing ADHB1^+^ CAFs and MYH11^+^ perivascular cells (**Fig. 4b**). Furthermore, spatial transcriptomic analysis of NSCLC revealed that *CCL19*-expressing perivascular cells and ADH1B^+^ CAFs were enriched in proximity to mature DC within TLS (**Extended data Fig. 4a-d**). Since CCR7 is the main receptor for CCL19, we examined whether deletion of CCR7 in cDC1 after T cells are primed and recruited to tumor sites compromises TLS maintenance in tumor tissues. To enable the temporal deletion of CCR7 specifically in the cDC1 subset, we reconstituted CD45.1^+^ mice with 1:1 (*Xcr1-DTR* CD45.2^+^:*Ccr7^-/-^* CD45.1^+^/CD45.2^+^) mixed bone marrow cells (**Fig. 4c**). DT administration at day 8 post tumor engraftment resulted in the depletion of the entire CCR7 wild type cDC1 compartment, whilst the remaining cDC1 lacked CCR7 expression **(Extended data Fig. 5a)**. Strikingly, CCR7 deletion from cDC1 at day 8 following tumor implantation significantly reduced the number and size of TLS and reduced the number of T_FH_ cells in lung tumors **(Fig. 4d)**. Similar findings were obtained upon CCR7 deletion from the whole cDC compartment using mice reconstituted with 1:1 (*Zbtb46-DTR* CD45.2^+^:*Ccr7^-/-^* CD45.1^+^/CD45.2^+^) mixed bone marrow (**Extended data Fig. 5a,b)**. Mature DC also express *CCL19* (**Fig. 4a**) however, conditional deletion of CCL19 from cDC1 did not alter TLS maintenance in tumors (**Fig. 4e**). These data establish that cDC1 migration to CCR7 ligand-enriched stromal hubs are required for TLS maintenance in tumor lesions.

A major role of mature cDC1 is to present tissue-associated antigens to T cells. Within the DC compartment, cDC1 are known to excel in the cross presentation of cell-associated antigens to both CD4^+^ and CD8^+^ T cells in secondary lymphoid organs (Eickhoff et al., 2015; Hor et al., 2015). cDC1 were also shown to provide a key scaffold to recruit CD4^+^ and CD8^+^ T cells enabling CD4^+^ T cells to provide helper cytokines in the vicinity of CD8^+^ T cells, which promotes their differentiation into CD8^+^ effector T cells in a process that requires cDC1 CD40 signaling (Eickhoff et al., 2015; Ferris et al., 2020; Hor et al., 2015). To investigate whether cDC1 cross-priming of CD4^+^ T cells was required for TLS maintenance in tumors, we reconstituted mice with 1:1 (*Xcr1- DTR* :*Cd40^-/-^*) mixed bone marrow cells to enable temporal depletion of CD40^+^ cDC1 upon DT administration, with the remaining cDC1 lacking CD40. DT administration 8 days post tumor implantation reduced TLS number and size, as well as the number of CD8^+^ T cells **(Extended data Fig. 5c)**.

To assess whether cDC1 sustained antigen presentation to CD4^+^ T cells was required to maintain TLS locally, we reconstituted mice with mixed bone marrow cells from *Xcr1-DTR* and MHC-II-KO mice and treated them with DT at day 8 post implantation. Strikingly, MHC-II deletion from cDC1 8 days after tumor implantation significantly reduced the number and size of TLS resulting in the dispersion of T cells and B cells within the tissue (**Fig. 4f**). Interestingly, this also reduced the number of T_FH,_ CD8^+^ T cells, and PD-1^+^ CD8^+^ T cells, increased the relative frequency of Tpex cells within this compartment, and reduced tumor-binding IgG antibodies (**Fig. 4g, h, Extended data Fig. 5d**). Using the same strategy, we also abrogated MHC-I cross-presentation of tumor antigens through conditional deletion of *B2m* from cDC1. Deletion of MHC-I^+^ cDC1 significantly reduced the number and size of TLS, albeit to a lesser extent than deletion of MHC-II^+^ cDC1 **(Fig. 4g)**. Disruption of MHC-I cross presentation did not affect the proportion of T_FH_ (**Fig. 4g**) or tumor-binding IgG antibodies (**Fig. 4g**). However, similar to the results observed upon MHC-II deletion from cDC1, deletion of MHC-I antigen presentation from cDC1 reduced the relative number of CD8^+^ T cells and PD-1^+^ CD8^+^ T cells and increased the relative frequency of Tpex cells within this compartment **(Fig. 4g, Extended data Fig. 5d)**. Interestingly, deletion of MHC-I or MHC-II from cDC1 resulted in a reduction of cDC1, suggesting that interactions with CD4^+^ and CD8^+^ T cells are required for their maintenance in tissue (**Fig. 4g**). Indeed, we observed that within TLS in human NSCLC, T cells expressed high levels of *FLT3LG* (**Fig. 4a**).

Interestingly, conditional deletion of IL-12 or IFNγR from cDC1 did not alter TLS maintenance **(Extended data Fig. 5e, f),** suggesting that intact IFNγR signaling in cDC1 is critical for efficient priming of T cells in the tdLN and the formation of TLS, but it is dispensable for the maintenance of TLS in tumors. In line with these results, mature DC within TLS in human NSCLC and HCC tumor lesions showed a downregulation of interferon-stimulated genes (ISG) compared to cDC1 (**Extended data Fig. 5g**), suggesting that IFNγR signaling does not control DC-T cell interactions in TLS once DC mature.

Taken together, our results reveal that *in situ* cDC1 migration to stromal hubs producing CCR7 ligands, and sustained presentation of tumor antigens to CD4^+^ and CD8^+^ T cells at the tumor site control the quantity and quality of TLS and enable the maintenance of T_FH_, the production of tumor-specific IgG antibodies and the differentiation of Tpex.

## Discussion

By leveraging spatial transcriptomics and multiplex imaging analysis of various human tumors and a novel murine model of NSCLC that forms mature TLS, we identified a critical role for the DC subset, cDC1, in the formation, maintenance, and function of TLS. Here we show that within the first week following tumor implantation, TLS formation relied on IFNγ-driven maturation of cDC1, their migration to the tdLN, and the subsequent recruitment of activated T cells to the tumor site. As tumors progressed, TLS maintenance became independent of tdLN T cell egress and instead depended on cDC1 migration to CCR7 ligand-enriched stromal hubs, sustained antigen presentation to intratumoral CD4^+^ and CD8^+^ T cells and CD40 signaling. These interactions were crucial for the maintenance of the T_FH_ pool, the formation of germinal centers and the production of tumor-specific antibodies, collectively sustaining TLS functionality and adaptive immune responses within the tumor microenvironment.

These results extend previous research indicating that DC present antigens to tumor-specific T cells within tumor-associated TLS (Joshi et al., 2015) and that DC control TLS maintenance in homeostatic conditions in the gut (McDonald et al., 2010) and in virally infected lungs (GeurtsvanKessel et al., 2009; Halle et al., 2009).

cDC1 ability to present antigens to both CD4^+^ and CD8^+^ T cells and to be licensed by CD4 T cells through CD40 signaling for cross-priming of CD8^+^ T cells in tdLN has been shown to be required for optimal anti-tumor immunity (Ferris et al., 2020) and anti-viral immune response (Eickhoff et al., 2015; Hor et al., 2015). By depleting cDC1 at a stage when TLS are no longer reliant on T cell recruitment from the tdLN, we demonstrate that cDC1-mediated antigen presentation to intratumoral CD4^+^ and CD8^+^ T cells, along with CD40 signaling, is essential for maintaining TLS structure and functionality. Much like in the LN, DC within TLS are strategically positioned in the T cell zone or at the interface between the T and B zones. Interestingly, we found that T_FH_ cells are present in TLS prior to germinal center formation, while depletion of cDC1 significantly reduced germinal center B cells in TLS, suggesting that interactions between mature DC and T_FH_ cells contribute to the germinal center response. In addition, similar to chronic viral infection where cDC1 preserve Tpex cells through MHC-I-dependent interactions within splenic niches (Dähling et al., 2022), we found that depletion of MHC-II and MHC-I from cDC1 promoted the accumulation of Tpex, suggesting that cDC1 within tumor-associated TLS contribute to Tpex differentiation into effector CD8^+^ T cells through direct MHC-I interaction and by offering proximal CD4^+^ help through MHC-II antigen presentation. This physical interaction may promote T_FH_ help to Tpex including through IL-21 as previously suggested (Cui et al., 2021; Magen et al., 2023). However, whether cognate interactions between cDC1, CD4^+^, and CD8^+^ T cells occur simultaneously or are distributed across distinct spatiotemporal niches remains to be determined. The importance of MHC-II presentation in anti-tumor immunity corroborates evidence that ICB response relies on both MHC class I and II presentation of tumor neoantigens in preclinical tumor models (Alspach et al., 2019) and that DC ability to present antigens to both CD4^+^ and CD8^+^ T cells are required for efficient tumor response to adoptive T cell therapy (Espinosa-Carrasco et al., 2024). Furthermore, clonal expansion of T_FH_ and effector CD8^+^ T cells strongly correlate with ICB response in patients with HCC (Magen et al., 2023). Similarly, our finding that alteration of TLS maintenance affect tumor antibodies formation reinforces the importance of TLS in the production of tumor-specific antibodies (Meylan et al., 2022; Ng et al., 2023).

We observed that TLS-associated DCs exhibit a distinct molecularly mature state, referred to as "mregDC" (Maier et al., 2020). The mregDC program is driven by tumor debris uptake and cholesterol mobilization (Belabed et al., 2023) and led to expression of CCR7 enabling the migration of mature DC to CCR7 ligand-enriched sites and efficient T cell priming. The enrichment of mature DCs in TLS suggests that antigen capture is a prerequisite for their migration and accumulation. We further identified CCL19-producing fibroblasts and perivascular cells as potential pivotal TLS organizers forming chemokine-rich hubs that guide the recruitment and interaction of CCR7^+^ mature DC, T cells, and B cells (Schumann et al., 2010). Interestingly, fibroblast reticular cells producing CCL19 were recently shown to promote T cell aggregates including TLS in NSCLC (Onder et al., 2024), and CRC liver metastasis (Zhang et al., 2024). Chronic microbial exposure in the lungs promotes CCL19 in stromal cells and drive the formation of TLS in the lung (Rangel-Moreno et al., 2011) and similar CCL19 hubs have been observed in chronic inflammatory skin lesions, psoriasis (Mitsui et al., 2012). These observations emphasize the potential key role of CCL19 in the formation of TLS, although the distinct contributions of CCL19 and the other CCR7 ligand CCL21 to TLS maintenance in tumors remains to be addressed. In addition to stromal hubs, DC and T_FH_ are also likely key organizers of TLS (Chaurio et al., 2022; Magen et al., 2023). Mature DC produce CCL17 and CCL22, two chemokine ligands known to recruit CCR4^+^ CD4^+^ T cells (Imai et al., 1999; Maier et al., 2020), while T_FH_ cells express high levels CH25h an enzyme that generate oxysterol gradients known to recruit Ebi2^+^ cells (Hannedouche et al., 2011), expressed on both DC and Tpex (Magen et al., 2023). Furthermore, cDC1 uniquely express XCR1 chemokine receptor, while its ligand, XCL1, is produced by Tpex cells (Im et al., 2016). The exact contribution of these molecules to TLS dynamics remains to be examined.

In summary, our study identifies cDC1 and their sustained antigen presentation to both CD4^+^ and CD8^+^ T cells as a pivotal driver of TLS formation and maintenance within tumors, emphasizing the importance of cDC1 in preserving TLS functionality. The interplay between TLS and tdLN in mediating tumor-specific immunity likely shifts throughout tumor progression, with TLS emerging as key contributors in more advanced tumors. In this setting, TLS may provide essential hubs to support the survival of antigen experienced lymphocytes and to amplify the efficiency of adaptive immune responses against tumor antigens. Our findings provide a foundation for novel therapeutic strategies aimed at enhancing effective antitumor T cell and B cell responses in cancer patients.

## Supporting information

Supplemental-Table-1-Patient_Clinical_Metadata

Supplemental-Table-2-MERFISH_Gene_Panels

Supplemental-Table-3-Antibodies

Supplemental-Table-4-Gene_List_Bulk_Seq

Supplemental-Table-5-Flow_Cytometry_Gating_Strategy

## Acknowledgments

We thank members of the Merad and Kamphorst laboratories at the Marc and Jennifer Lipschultz Precision Immunology Institute at Mount Sinai for insightful discussions and feedback; and the Mount Sinai Flow Cytometry Core, the Microscopy and Advanced Bioimaging CoRE, the Human Immune Monitoring Center (HIMC), the Center for Comparative Medicine and Surgery for animal husbandry and Biorepository and Pathology CoRE Laboratory at the Icahn School of Medicine at Mount Sinai for support. We thank the patients and their families for participating in the clinical studies. We recognize the invaluable assistance of Barzin Y. Nabet (Genentech), Alfonso Rodriguez Sanchez-Paulete, John A. Grout and Mark Buckup (Icahn School of Medicine at Mount Sinai). Schematic figures have been created with BioRender under academic licence. This work was supported in part through the computational resources and staff expertise provided by Scientific Computing at the Icahn School of Medicine at Mount Sinai. We are grateful to Boehringer Ingelheim Inc. for their sponsorship and support in providing resources for CRC sample collection. R. Mattiuz was supported by the 2021 AACR-AstraZeneca Immuno-oncology Research Fellowship, Grant Number 21-40-12-MATT and by the Portuguese Foundation for Science and Technology (grant number 2023.15874.PEX). M.D.P. was supported by the American Federation for Aging Research (2021 Diana Jacobs Kalman Scholarship for Research in the Biology of Aging). S.H. was supported by the National Cancer Institute predoctoral-to-postdoctoral fellowship (K00 CA223043). S.K.S. was supported by grants from the National Institutes of Health (U24 CA224319, R01 DK1222853, U2C ES030859, DA055434, 1R01 HL166720-01A1, U01OH012621) and funding from Caribou Biosciences, Inc. S. Gnjatic was partially supported by National Institutes of Health (NIH) grants CA224319, DK124165, CA263705 and CA196521. A.O.K. and T.U.M. were supported in part by the Tisch Cancer Institute Cancer Center Support Grant (P30 CA196521). A.O.K. was supported in part by R01 AI153363. M.M. was partially supported by NIH grants CA257195, CA254104 and CA154947. A.L. was supported by NIH/NCI R37CA230636, NIH/NCI R01CA251155, and Damon Runyon-Rachleff Innovator Award.

## Author contributions

R. Mattiuz and M.M. conceptualized and obtained funding for the project. R. Mattiuz and J.LB. performed experiments with the help of J.B., P.H., A.V., M.D.P., M.B., E.H., R. Merand, S.H. and E.N. B.Y.S., L.H., E.R., M.D.P., D.D., I.F., E.G.K., V.R. performed the computational analysis. A.V. and R.D. modified the KP-HELLO cell line. R. Mattiuz, P.H., A.V., H.M.K., L.T., M.K., I.P.M., G.I., S.O., A. Tepper, H.M., E.N., S. Goldstein, A.M.R., A Tabachnikova and H.S. performed multiplex imaging experiments and analyzes. M.B. and G.M. did the spheroid experiments. R. Mattiuz, P.H., L.T. and S.H. processed the human samples. CH coordinated the clinical and research teams and manage clinical specimens. S.O. and A.D.P. provided pathological expertise for tissue annotation. M.N., J.A.K., N.B., A.L., Z.C., S.K.S., J.D.B., M.S., C.M., S. Gnjatic, C.S.F., W.H.F., B.D.B., T.U.M., J.G.C., H.S., T.C.B., N.S.J. and A.O.K. provided intellectual input, essential reagents and datasets. R. Mattiuz wrote the original draft. R. Mattiuz, J.B. and M.M wrote, reviewed and edited the manuscript. All authors provided feedback on the manuscript draft.

## Material and methods

### Mouse strains

C57BL/6, B6.Cg-Zbtb46tm3.1(cre)Mnz/J (strain #028538), B6.129P2(C)-Ccr7tm1Rfor/J (#006621), C57BL/6N-Ifngr1tm1.1Rds/J (#025394), B6.129S(C)-Batf3tm1Kmm/J (#013755), C57BL/6-Gt(ROSA)26Sortm1(HBEGF)Awai/J (#007900), B6(129S4)-Xcr1tm1.1(cre)Kmm/J (#035435), SMARTA (H2-I-Ab-restricted LCMV GP61-80 epitope) CD45.1 (#030450) mice were purchased from JAX and *Ccr7* floxed mice were kindly donated by Dr. Milena Bogunovic and Dr. Iannis Aifantis. For bone marrow transplant experiments, CD45.1, B6.129P2-Cd40tm1Kik/J (strain#002928), B6.129P2-B2mtm1Unc/DcrJ (#002087), B6.129S2-H2dlAb1-Ea/J (#003584), B6(Cg)-Zbtb46tm1(HBEGF)Mnz/J (#019506), B6.129S1-Il12btm1Jm/J (#002693) mice were also purchased from JAX and *Ccl19^-/-^* mice were kindly donated by Dr. Jason G. Cyster. All animal experiments performed in this study were approved by the Institutional Animal Care and Use Committee at the Icahn School of Medicine at Mount Sinai. Mice within experiments were age and sex-matched. Tumor implantations and were conducted in mice between 7–15 weeks of age. Mice were housed in individually ventilated cages at the Mount Sinai specific-pathogen-free (SPF) facilities, provided food and water ad libitum, with conditions maintained at 21∼23C and 39∼50% humidity and 12/12 hour dark/light cycle.

### Bone marrow transplantation

Bone marrow (BM) chimeras were generated by retro-orbitally injecting 1-10x10^6^ total donor cells (fresh or cryopreserved BM) into sub-lethally irradiated 6-week-old recipient mice (two doses of 5.5 Gy administered 6 hours apart). A period of 9 to 12 weeks was granted to ensure engraftment. Recipients were supplemented with sulfamethoxazole/trimethoprim for 3 weeks. At 9 weeks, successful reconstitution (minimum 90%) was assessed by flow cytometry analysis of peripheral blood.

### Generation of KP-HELLO-2 tumor model

KP-HELLO-2 cells were originally derived from the KP-HELLO cells (kindly provided by Dr. Nikhil S. Joshi, (Cui et al., 2021)) and modified in house. MSCV-IRES-Thy1.1 DEST was a gift from Anjana Rao (Addgene Cat # 17442), and gamma-retrovirus encoding Thy1.1 was produced by transfecting HEK 293T cells with Thy1.1 vector along with gamma retroviral packaging plasmids. Briefly, KP-HELLO cells were expanded on a tissue culture-coated plate. Upon reaching 60% confluence, they were transduced with retroviral particles by spinfection with polybrene (fiinal concentration of 8µg/ml). Cells were expanded in RPMI supplemented with 10% FBS and 1% Pen-strep, Glutamax and HEPES and Thy1.1^+^ cells (stained with anti-Thy-1.1 PE-Cy7, clone: OX-7, BioLegend Cat# 202518) were sorted twice over 10 passages on a CytoFLEX SRT Cell Sorter (Beckman).

### Cell lines

KP-HELLO-2, KP-HELLO and KPAR1.3 (kindly provided by Dr. Julian Downward, (Boumelha et al., 2022)), cells were derived from a *Kras*^LSL-G12D/+^;*p53*^fl/fl^ background, AKPS colorectal tumor organoids from a *Apc*^fl/fl^;*Kras*^LSL-G12D/+^;*p53*^fl/fl^;*Smad4*^fl/fl^ background (kindly provided by Dr. Ömer H. Yilmaz, (Goto et al., 2024)) and Hep-53.4 HCC cells from a carcinogen-induced liver model (Kress et al., 1992); Cellosaurus CVCL_5765) were kindly provided by Dr. Josep Llovet and originally obtained from Cytion (Product number 400200). All cell lines were grown in in complete cell culture medium (DMEM + 10% FBS + 1% P/S).

### Orthotopic tumor models

For orthotopic lung tumors, 1.5 x 10^5^ KP-HELLO-2, KP-HELLO or KPAR1.3 cells were intravenously injected into the tail vein. Tumor-bearing lungs and tumor-draining lymph nodes were analyzed 15 days post-injection, unless otherwise specified in the figure legends. For KPAR1.3, tumors were harvested 21 days post-injection. To assess tumor burden, the left lung lobe was fixed in paraformaldehyde, embedded in paraffin, and examined as 5 µm cross-sections. Upon hematoxylin and eosin staining, lung tissue sections were scanned on slides using an Olympus digital scanner and analyzed using the Panoramic viewer and QuPath software. For orthotopic CRC tumors, mice received intra-ceacum injection of 1.5 x 10^5^ AKPS cells in 10 uL basement-membrane extract (BME, R&D Systems Cat # 35-330-0502) and tumors were analyzed six weeks post-injection. For orthotopic HCC tumors, mice were injected orthotopically into the left-lateral lobe of the liver with 1 x 10^6^ Hep53-4 cells in 10ul PBS and tumors were harvested 2 weeks post-injection.

### *In vivo* treatments

For prolonged and effective conditional depletion of cDCs or cDC1, *Zbtb46-hDTR* and *Xcr1^Cre/wt^; Rosa26^hDTR/wt^* mice received an initial dose of DT (32 ng per g body weight, List Biological Laboratories Cat # 150), followed by injections of 20 ng per g every 60 hours. To inhibit lymphoid cell egress from peripheral lymphoid organs, mice were administered 20 µg of FTY720 (Cayman Chemical, Cat # 10006292) daily over the indicated time periods. To increase DC numbers, mice received 75 µg of FLT3-L-Fc (Gilead) on day 8 post-tumor engraftment.

### Adoptive cell transfer

In experiments analyzing tumor-specific CD4 T cells (SMARTA; CD4^+^ T cells specific for the LCMV GP66–77 epitope), CD4^+^ T cells were isolated from blood of naïve SMARTA CD45.1 transgenic mice with the EasySep Mouse CD4^+^ T Cell Isolation Kit (StemCell, Cat # 19852). Between 20-50 x 10^3^ naïve SMARTA CD4^+^ T cells were transferred intravenously into C57BL/6 recipient mice one day before tumor injection.

### Flow cytometry

Single-cell suspensions were obtained from tumor bearing lung and tumor draining lymph nodes by digestion with 0.25 mg/ml collagenase IV (Sigma, Cat # C5138-1G) at 37 °C for 30 min (lung) or 25 min (lymph nodes) followed by passing through a 70-μm cell strainer and red blood cell lysis (RBC lysis buffer, BioLegend, Cat # 420301) for 2 min at room temperature (RT). For T cell cytokine assessment, cells were incubated with 1 μg/ml brefeldin A, 1 μg/ml ionomycin, and 50 ng/ml PMA (all from Sigma, Cat # B7651, I0634, P1585) for 4 hours at 37 °C. Cells were stained in FACS buffer (PBS supplemented with 10% BSA and 2 mM EDTA) for 25 min at 4°C. For CCR7 and CXCR5 staining, cells were incubated for 30 min at 37°C. To assess tumor-specific CD4^+^ T cells, cells were stained at 37°C for 2 hours with an APC-conjugated MHC-II LCMV Gp66 tetramer (produced by the NIH Tetramer Core Facility and kindly provided by Dr. Alice O. Kamphorst) used at a 1:100 dilution. Cytokines and transcription factors were stained following fixation and permeabilization using the Foxp3/Transcription Factor Staining Buffer Set (eBioscience, Cat # 00-5523-00). Samples Cells were analyzed with BD LSR Fortessa or BD FACSymphony analyzers (BD Biosciences). Flow cytometry data was acquired using FACS Diva software v.9 (BD), and the data obtained were analyzed using FlowJo (LLC). Antibodies used are listed in Supplemental Table 3. Gating strategies are shown in Supplemental Table 5.

### Tumor-specific antibody binding assay

To measure tumor-binding antibodies, serum was collected from tumor-bearing mice at end-point and heat-inactivated at 56 °C for 10 min. KP-HELLO-2 cells were stained with serum (1:50) for 40 minutes at 4 °C followed by a secondary staining with anti-mouse IgG (1:200, Poly4053-PE- Cy7, Biolegend, Cat # 405315) or anti-mouse IgM (1:200, II/41-PerCP-eFluor710, eBioscience, Cat # 46-5790-82) for 40 minutes at 4 °C, as previously described (Ng et al., 2023).

### Immune aggregate formation using KP-HELLO-2 spheroids co-cultured with mutuDCs and TILs

KP-HELLO-2 spheroids were cultured for 7 days as previously described (Lugand et al., 2022) with 10,000 KP-HELLO-2 cells expressing mScarlet per spheroid, using the Spherotribe kit (Idylle). In parallel, mutuDCs were cultured with KP-HELLO-2 cell debris to induce maturation which was confirmed by flow cytometry (Belabed et al., 2023; Fuertes Marraco et al., 2015). Separately, TILs were harvested and sorted from KP-HELLO-2 tumors 7 days post-injection, maintaining the endogenous T cell/B cell ratio, and labeled with DeepRed cell tracker for 30 minutes. Mature mutuDCs were added to KP-HELLO-2 spheroids for 4 hours, followed by the addition of TILs (30,000 mutuDCs and 30,000 TILs per spheroid). After 5 days, KP-HELLO-2 spheroid size and subsequent immune attraction were assessed by microscopy and ImageJ. Spheroid infiltration by TILs and 3D aggregate formation in the presence of mutuDCs were measured by BiPhoton microscopy and analyzed with QuPath. The total number of TILs in KP-HELLO-2 spheroids co-cultured with mutuDCs was also measured by flow cytometry

### Immunofluorescence imaging for TLS assessment

5um FFPE slides were baked at 60 °C for 2 hours before deparaffinization. Slides were deparaffinized in xylene (two treatments) and rehydrated through a series of ethanol solutions of decreasing concentrations (from 100% to 70%), followed by PBS rinses. For antigen retrieval, slides were immersed in pre-warmed DAKO pH 9 solution (with or without 10% glycerol) at 95°C, then allowed to cool to room temperature and washed in PBS. Blocking was performed in a humidified chamber with a blocking buffer (1X TBS, 10% BSA, 0.1% Triton X-100). Autofluorescence was quenched with TrueBlack solution, followed by a PBS wash. Slides were incubated with primary antibody (1:200) in blocking buffer overnight at 4°C. After PBS washes, slides were incubated with secondary antibody, diluted 1:500 in PBS and containing 2% serum from the host species, and counterstained with DAPI. Finally, slides were mounted with ProLong Gold Antifade Mountant (Invitrogen, Cat # P36930), coverslipped, and stored at 4°C. Whole stained slides were scanned on a CyteFinder HT II fluorescence scanner (RareCyte) equipped with a 20x lens. Images were corrected for optical distortion, stitched and processed locally using the on-board CyteHub software. Alternatively, whole-slide images were captured using a Leica DMi8 microscope. Antibodies used are listed in Supplemental Table 3.

### Machine learning TLS detection using Qupath

To detect TLS at a whole-slide level, Random Trees pixel classifiers were applied in QuPath (Bankhead et al., 2017) with a resolution setting of 5.20 µm/pixel, selecting three channels (Alexa 488, Alexa 647, and DAPI) at scales of 2.0 and 4.0, using Gaussian features without normalization. Classifiers were trained on at least five manually annotated TLS regions (aggregates of B220^+^ cells, CD3^+^ cells, and DAPI^+^ cells) and five non-TLS regions. The output classifications were then segmented into individual annotations with areas exceeding 1500 µm². Immune aggregate classifications were reviewed and validated by a trained pathologist.

### Tumor antigen-specific B and T cell imaging

Prior to immunofluorescence staining, 10 µm OCT slides were fixed in acetone at -20°C for 10- 15 minutes and dried. To detect tumor-antigen-specific B cells targeting hen egg lysozyme (HEL), HEL protein (Sigma, Cat # 10837059001) was conjugated to Alexa Fluor 488 using the Alexa Fluor 488 Protein Labeling Kit (ThermoFisher) according to the 20-50 kDa protein protocol. OCT-embedded slides were stained with HEL-Alexa488 (1:500) and co-stained with B220, GL7, and DAPI. To detect tumor-antigen-specific CD4^+^ T cells recognizing the LCMV GP66–77 epitope, OCT-embedded slides from adoptively transferred SMARTA CD45.1 mice were stained with CD45.1 and co-stained with CD4, PD-1, and DAPI (Supplemental Table 3).

### Multispectral imaging for TLS maturity assessment

For multispectral staining, every cycle of antigen staining was processed as follows. The tissues were subjected to HIER either in AR6 or AR9 citrate buffers (Akoya Biosciences). Post-antigen retrieval, slides were blocked with 1X antibody diluent/block (Akoya Biosciences) for 10 minutes, followed by staining with primary antibody for 30 minutes. Every staining step took place in a humidified chamber at room temperature. After the wash with 1X TBS-Tween (TBST) buffer (3 times 2 minutes each), the slides were stained with HRP-conjugated secondary antibody, either anti-rabbit IgG (Fisher, Cat # MP740150) or anti-Rat IgG (Biocare, Cat # RT517L) for 30 minutes. Then after the wash with 1X TBST, separate opal detector fluorophores were used for each marker. TLS maturity panel was as follows: CD4 (1:200)/Opal 480, CD8 (1:400)/Opal 780, CD19 (1:1000)/Opal 520, Ki67 (Clone SP6, ThermoFisher, 1:100, Cat # MA5-14520)/Opal 570, CD21 (Clone SP186, Abcam, 1:50, Cat # ab227662)/Opal 620, and AID (Clone mAID-2, ThermoFisher, 1:800, Cat # 14-5959-82)/Opal 690. The last round of staining was carried out with additional antigen retrieval and DAPI nuclear staining. The staining was performed either manually or on automated machine (Opal 6-Plex Detection Kit, Akoya Biosciences, Cat # NEL871001KT). Then the stained tissue sections were mounted with Diamond Anti-fade mounting media and were imaged as whole slides scans on the Vectra (Perkin Elmer). Images were unmixed and were analyzed on QuPath.

### QuPath Tissue Analysis for TLS maturity assessment

QuPath software was used for quantification analysis. Cell detection and segmentation was carried out using StarDist extension based on DAPI nuclear stains (Schmidt et al., 2018) For phenotyping of individual cells, machine learning approach was utilized to generate an object classifier for each marker. In the case of markers that are mutually exclusive, we generated a group object classifier that includes all those markers. Then, these individual classifiers were compiled together and applied onto multispectral images to phenotype cells. For annotations of TLS, machine learning approach was once again used to generate a pixel classification with the annotation size greater than 1500 um^2^ (MacFawn et al., 2024; Ruffin et al., 2021).

### Cyclic immunofluorescence (CyCIF)

FFPE sections were prepared and stained using an 18-plex antibody panel following previously described CyCIF methodologies. For sample preparation, slides were automatically baked at 60 °C for 30 minutes and dewaxed at 72 °C using BOND Dewax Solution. Antigen retrieval was carried out at 100 °C for 20 minutes in BOND Epitope Retrieval Solution 2 (ER2) using a Leica Bond RX system. To minimize autofluorescence, slides were treated with a bleaching solution (4.5% H2O2, 20 mM NaOH in PBS) and exposed to LED light for 2 cycles of 45 minutes. To reduce non-specific antibody interactions, slides were rinsed in PBS for 3 x 5 minutes and incubated overnight at 4 °C with secondary antibodies diluted 1:1000 in 150 µL of Odyssey Blocking Buffer, protected from light. Slides were then washed 3 times with PBS and bleached again for 2 x 45 minutes. For each round of CyCIF, samples were incubated overnight at 4 °C in the dark with Hoechst 33342 (1:10,000; Thermo Fisher Scientific, Cat # H3570) for nuclear staining, alongside either primary conjugated antibodies or primary unconjugated antibodies diluted as specified (Supplemental Table 3) in 150 µL of Odyssey Blocking Buffer (LI-Cor). When using primary unconjugated antibodies, incubation was followed by a 2-hour room temperature incubation with secondary antibodies in the dark. Subsequently, slides were washed 3 x 5 minutes and mounted with 200 µL of 70% glycerol. Imaging was performed automatically on a RareCyte Cytefinder II HT system, with exposure settings optimized per channel to prevent signal saturation while remaining consistent across samples. After imaging, coverslips were removed by placing slides in 1X PBS and heating them in a water bath for 1 hour. Between cycles, slides were photobleached for 2 x 45 minutes and washed 3 x 5 minutes in PBS (Lin et al., 2023, 2018; Mollaoglu et al., 2024).

### CyCIF image preprocessing and quality control

The entire pre-analytical CyCIF image processing workflow, including stitching, registration, illumination correction, segmentation, and single-cell feature extraction, was carried out using the MCMICRO pipeline, an open-source multiple-choice microscopy platform (full code available at GitHub: https://github.com/labsyspharm/mcmicro). For probability map generation, a pre-trained U-Net model, UnMicst v2, was applied, followed by a marker-controlled watershed algorithm for single-cell segmentation. Nuclear detection was performed within a diameter range of 3 to 60 pixels. Probability maps generated by UnMicst were subsequently processed with S3segmenter to produce nuclear segmentation masks, while the cytoplasmic regions were defined by expanding the nuclear mask by 3 pixels. After segmentation masks were created, mean fluorescence intensities for each marker were computed for each cell, resulting in a single-cell data table corresponding to each acquired whole-slide CyCIF image. Annotated histologic regions on the whole-slide image were used to extract quantified single-cell data for cells located within the specified ROI range. Multiple strategies were employed to ensure the accuracy and reliability of the single-cell data. At the image level, cross-cycle image registration and tissue integrity were reviewed, and poorly registered regions or areas with deformed tissues or artifacts were identified and excluded from the analysis. Antibodies exhibiting low-confidence staining patterns, as determined through visual inspection, were also removed from the analysis. Segmentation quality was rigorously assessed, and segmentation parameters were refined iteratively to enhance the precision of the segmentation masks. At the single-cell data level, correlations of DNA staining intensities across different cycles were evaluated to filter out cells lost during the cyclic process, using a correlation coefficient threshold of less than 0.8 for exclusion (Schapiro et al., 2022).

### Multiplex IHC imaging

FFPE sections (4µm) were stained using the multiplexed immunohistochemical consecutive staining on a single slide (MICSSS) protocol as previously described (Remark et al., 2016). Briefly, slides were baked at 50°C overnight, deparaffinized in xylene and rehydrated in decreasing concentration of ethanol (100%, 90%, 70%, 50% and dH2O). Sample slides were incubated in pH6 or pH9 buffers at 95°C for 30min for antigen retrieval, then in 3% hydrogen peroxide for 15min and in serum-free protein block solution (Dako, Cat # X090930-2) for 30min. Primary antibody staining was performed using the optimized dilution during 1h at room temperature or at 4°C overnight followed by signal amplification using associated secondary antibody conjugated to horseradish peroxidase during 30min. Chromogenic revelation was performed using AEC (Vector, Cat # SK-4200). Tissue sections were counterstained with hematoxylin, mounted with a glycerol-based mounting medium and finally scanned to obtain digital images (Aperio AT2, Leica). After scanning, slide coverslips were removed in hot water (∼50°C). Primary antibodies are presented in Supplemental Table 3.

To quantitatively analyze localization and co-expression patterns, tissue annotation was first done by pathologists using QuPath software. For the analysis, we used the svs multi-resolution, pyramidal images obtained per marker after staining. Both the AEC chromogen stain and the hematoxylin nuclear counterstain were extracted from each image via a dynamically determined deconvolution matrix. Then, each image was split into smaller tiles to permit computational analysis. Each tile from the first stained image was matched to the respective tile from the sequential stained images and then elastically registered using the extracted hematoxylin nuclear stain and SimpleElastix open-source software. Then, by using an iterative nuclear masking via STARDIST, we produced a composite semantic segmentation for nuclei residing in the series of tiles. Each nucleus was artificially expanded by a number of pixels to simulate a cytoplasm per cell and that was coherent with membrane marker staining. Lastly, cellular-resolution metadata was acquired for all cells in the final cell mask. To unbiasedly determine positive and negative cells per marker, we used an unsupervised classification technique to cluster cell populations, followed by a supervised approach where we would evaluate each cluster as positive or negative per marker. First, the metadata aggregated per sample was collected, transformed to z-scores, randomized, and split into subsamples per batch. Each batch was processed in parallel: data for each marker was transformed, clustered and collapsed into multiple groups by principal components analysis and uniform manifold approximation and projection. Then, we performed the final quality control to manually attribute which clusters were positive or negative. This produced a final cellular-resolution data frame containing binary marker classification that was used for downstream localization, marker co-expression, tissue annotation and reconciliation and statistical analyses.

### Human subjects

Samples of tumor were obtained from patients undergoing surgical resection at Mount Sinai Hospital (New York, NY) after obtaining informed consent in accordance with a protocol reviewed and approved by the Institutional Review Board at the Icahn School of Medicine at Mount Sinai (IRB Human Subjects Electronic Research Application 18-00407 for HCC samples, 10-00472 and 10-00135 for NSCLC samples and 18-00407 and 18-00855 for CRC samples) and in collaboration with the Biorepository and Department of Pathology. For multiplex imaging of patient TLS maturity conducted at the University of Pittsburgh, Pittsburgh, PA, the studies were performed under IRB protocols STUDY19060269 (NSCLC patients) and PRO17080326 (HGSOC patients).

The single-arm, open-label, phase 2 trial of HCC patients with resectable tumors was registered on ClinicalTrials.gov (NCT03916627, Cohort B). Twenty patients were enrolled and received two cycles of Cemiplimab before surgical resection as described in (Marron et al., 2022). Response to treatment was defined as more than 50% (partial response) or more than 70% (complete response) necrosis of resected tumor by pathologists. Similarly, TLSHi versus TLSLo was defined after annotation and scoring of H&E samples by the pathologist. Post-operatively patients underwent standard-of-care screening every 6 months; recurrence was defined as development of imaging consistent with recurrent HCC using the Liver Imaging Reporting and Data Systems (LIRADS) score of 5 for an intrahepatic lesion (the most common site of HCC recurrence) or arterial enhancing new extrahepatic metastatic foci consistent with metastatic disease (https://aasldpubs.onlinelibrary.wiley.com/doi/pdf/10.1002/hep.27304). The patient clinical metadata can be found in Supplemental Table 1.

### 10X Visium spatial transcriptomics

#### 10X Visium spatial sequencing library preparation

Prior to processing Visium gene expression samples, DV200 was measured on RNA isolated from Formalin-fixed paraffin-embedded (FFPE) blocks to assess RNA integrity. Samples with a DV200 greater than 50% were selected for subsequent steps. After quality control, blocks were trimmed and 5-µm tissue sections were cut and placed on the capture area of a Visium Spatial Gene Expression for FFPE slide (10X Genomics). The slide was incubated in a thermal cycler at 42 °C for 3 h, and then placed in a desiccator overnight at room temperature. Subsequent steps covering tissue staining and library preparation were performed according to the manufacturer’s instructions (CG000407_VisiumSpatialGeneExpressionforFFPE_RevC). The quality of the libraries was assessed using an Agilent 4200 TapeStation. Libraries were sequenced using the NovaSeq 6000 platform with 150 paired-end configuration on a NovaSeq SP flowcell.

#### Tested Visium FFPE samples with immune aggregates

The tested samples included in-house generated data for lung (n=3), liver (n=3) and colorectal (n=6) human cancers and third-party data set for renal (n=6; GSE175540) human cancer. All the tested samples were stained using hematoxylin and eosin. Visium transcriptome data was obtained using standard procedure provided by 10X Genomics for FFPE tissue sections. Reads were processed and mapped to the human reference genome *GRCh38* using the 10x Genomics Space Ranger pipeline (10X Genomics). Filtering and analysis of the samples were performed in python using scanpy (v1.9.1), with default parameters. In the downstream analysis, samples retained for further study were those containing at least one TLS per tissue area, as confirmed by a pathologist.

#### Deconvolution of Visium spots for spatial cell type mapping

For the deconvolution step we used Cell2location algorithm (Kleshchevnikov et al., 2022) yielding proportions of the deconvoluted cells in each Visium spot. We also performed hematoxylin and eosin (H&E)-based cell segmentation to quantify the number of cells per each Visium spot using StarDist (Schmidt et al., 2018). Then, we multiplied deconvoluted proportions of cells by the absolute number of segmented cells in each spot, thus getting the absolute number of deconvoluted cells in each Visium spot.

#### Spatial enrichment in TLS regions

For each Visium sample, we used the histopathological annotations of TLS as base to test the enrichment or depletion of inferred cell types. First, the inferred cell type abundance values are binarized based on their spatial autocorrelation using *Voyager* https://pachterlab.github.io/voyager/index.html, (Moses et al., 2023). Univariate local statistics are calculated with the *runUnivariate* function (Getis-Ord GI* with permutation testing; type = “localG_perm”) and spots were called if value > 2. Enrichment fold changes and statistics were calculated using *LotOfCells* (González-Velasco, 2024). Correlation and hierarchical trees were calculated by *cor* and *hclust* functions. Visualizations were generated in R.

#### Gradient analysis

Gradient analysis consists of two steps. First, it identifies Visium spots positioned equidistantly from a selected structural boundary, whether they are inside the structure (internal layers) or moving away from it (outer layers). Then, it regresses gene expression or cellular counts across these layers and thus can explore the microenvironment of a selected tissue region (Radkevich et al., 2024).

#### Radial distance analysis from TLS regions

For each Visium sample, we used the TLS annotations to extract the radial distances, *d*, using the *RadialDistance* function in *semla* (convert_to_microns = T; v1.1.6; cit) (Larsson et al., 2023). Distances of 0 µm represent the outer line of spots immediately exterior to the TLS-labeled spots. Distances >1,000 µm were discarded to exclude distant tissue areas. In *ggplot2*, *geom_smooth* with the locally estimated scatterplot smoothing method (“loess”, span = 0.45) was used for visualization of cell density data.

#### Pathway enrichment analysis of Visium samples

We used GSEApy (Fang et al., 2023) implementation of gene set enrichment analysis. Used databases of pathways – GO Cellular Component (https://geneontology.org/), KEGG (https://www.genome.jp/kegg/), LINCS (https://lincsproject.org/), MSigDB (https://www.gsea-msigdb.org/gsea/msigdb) and Reactome (https://reactome.org/).

#### Code sharing statement for analysis of Visium samples

All the code used for cell segmentation in Visium samples is available at the Human Immune Monitoring Center (HIMC) GitHub page https://github.com/ismms-himc/visium_segmentation.

The code for gradient analysis is deposited in the HIMC GitHub page, https://github.com/ismms-himc/Visium_analysis.

#### Other packages used for visium analysis

Seurat (Hao et al., 2024), Simple features (https://doi.org/10.32614/RJ-2018-009), Tidyverse (Wickham et al., 2019), Scran (Lun et al., 2016), Scater (McCarthy et al., 2017).

### scRNA-seq analysis

Previously published NSCLC and HCC scRNA-seq datasets (GSE154826, GSE206325) were analyzed. Differentially expressed genes were determined using Seurat. Imputed mean UMI counts were used to calculate logarithmic fold changes in expression between cell states to further analyze markers of interest. Gene set enrichment analysis was conducted using the Enrichr database. Additional R packages used include scDissector v1.0.0, shiny v1.7.0, ShinyTree v0.2.7, heatmaply v1.3.0, plotly v4.10.0, ggvis v0.4.7, ggplot2 v3.3.5, dplyr v1.0.7, Matrix v0.9.8, and seriation v1.3.5.

### MERFISH

To identify transcriptionally distinct cell population with MERFISH, we used two panels of genes as previously described (Magen et al., 2023), Supplemental Table 2. FFPE or fresh frozen samples from HCC and NSCLC patients were processed as previously described and tissue sections were incubated with a custom designed MERSCOPE Gene Panel Mix for 36-48 hours at 37°C. Samples were gel embedded and underwent tissue clearing as previously described. Tissue slides were prepared for imaging as previously described and loaded onto the MERSCOPE system (Vizgen 10000001). Samples were initially imaged at low-resolution using 10x magnification to select regions of interest which were imaged at high-resolution using 60x magnification.

Images were visualized using MERSCOPE Vizualizer (v2.1.2593.1) and single-cell analysis was conducted using the the scanpy (v1.9.1) Python package as previously described. Briefly, cells with <10 and >750 counts or with <10 unique genes expressed were removed. Furthermore, cells with a high fraction of blank counts (top 5^th^ percentile) or with high or low spot density or polyT signaling staining intensity (top and bottom 0.0005 percentile) were filtered to remove debris, apoptotic cells, and suspected doublets. Label transfer was performed using TACCO (v0.2.2) from annotated single-cell data to the MERFISH data using the MERFISH raw counts as inputs and default parameters for label transfer. For HCC samples, a previously published single-cell reference was used (Magen et al., 2023). For NSCLC samples, a mixture of the liver tumor single-cell reference and a prior lung adenocarcinoma atlas (Leader et al., 2021) was used due to the increased resolution of desired immune cell subtypes in the liver data. Specifically, clusters labeled as T, NK, DC, B memory, and B naive from the HCC dataset replaced respective cell types in the NSCLC dataset. Resulting labels were compared against scanpy-based Leiden clustering of MERFISH data for concordance of broad cell type compartments. After label transfer, tissue regions were drawn using Shapely (v2.0.1) and GeoPandas (v0.14.0) around cells labeled naive B and memory B cells, which were annotated as “immune aggregates”. In HCC samples, regions were drawn around tumor, endothelial, and hepatic stellate cells, which were annotated as “stromal”. In NSCLC sections, similar “stromal” regions were created around epithelial, endothelial, and fibroblast cell types. Regions were drawn as alpha shapes around cells, using the centroid x and y locations of each cell as inputs. To smooth regions of sparse cells, all regions were buffered by 30 microns. Overlapping areas of regions were resolved such that immune aggregate regions were prioritized first prior to stromal. Co-occurrence analysis was performed separately for cells of interest in stromal and immune aggregate regions using squidpy (v1.5.0) (Palla et al., 2022) using the squidpy.gr.co_occurrence() function at regular intervals from 0 to 500 microns and the TACCO assigned cell type labels as inputs. Enrichment values for each cell type at 30 microns were normalized for visualization by ln(1 + x), where x is the co-occurrence enrichment value as derived from squidpy. Co-occurrence results were also checked for agreement with permutation-based neighborhood enrichment, calculated through the squidpy.gr.nhood_enrichment() function. All MERFISH downstream processing and analysis was conducted on a distributed high performance computer cluster. This work was supported in part through the computational resources and staff expertise provided by Scientific Computing at the Icahn School of Medicine at Mount Sinai and supported by the Clinical and Translational Science Awards (CTSA) grant UL1TR004419 from the National Center for Advancing Translational Sciences.

### POPLAR cohort

*Survival analysis.* Overall survival analyses of the POPLAR cohort (Fehrenbacher et al., 2016; Mazieres et al., 2021; Patil et al., 2022) was performed using a Kaplan-Meier estimator and Cox regression upon stratifying patients by presence of tertiary lymphoid structures (TLS) or lymphoid aggregates (LA). TLS or LA annotations were determined by histological imaging. R packages used include: survival, survminer, and gtsummary.

### Immune cell scores and pathway analyses

Gene signatures for different immune cell types were generated from prior literature (Leader et al., 2021; Magen et al., 2023) cf. Supplemental Table 4. Scores for each signature were computed as the mean of z-scores for each gene across patient samples. To determine correlations between immune cell scores, Pearson correlation coefficients were computed (two-tailed, 95% confidence interval). These signatures were also used to determine significant cell signaling pathways via gene set enrichment analysis (Xie et al., 2021) (Xie et al., 2021).

## Data availability

No new software pipelines were used in the study beyond those described in relevant Methods sections and in (Radkevich et al., 2024). Accession numbers for reanalyzed published datasets include the human renal cancer FFPE Visium dataset (GSE175540), external NSCLC bulk RNA-seq dataset (EGAS00001005013), and NSCLC and HCC single-cell RNA sequencing (scRNA - seq) datasets (GSE154826, GSE206325, GSE183219). Processed matrix files and metadata for the human cancer FFPE Visium dataset and human HCC and NSCLC MERFISH datasets generated in this study will be publicly available at the time of publication. Any additional information required to interpret data reported in this paper is available from the corresponding author upon reasonable request.

### Statistics

Parametric statistical tests, such as the two-tailed Student’s t-test (for unpaired comparisons) and paired t-test (for paired comparisons), were used to assess differences between two groups when the data met normality assumptions. For non-parametric comparisons, normality was first evaluated using the Kolmogorov–Smirnov test (for unpaired data) or the Shapiro–Wilk test (for paired data). If the data significantly deviated from normality, a Mann–Whitney U test was used for unpaired comparisons, and a Wilcoxon signed-rank test was performed for paired samples. Across all panels, data represent mean ± SEM.

---

**Extended Fig 1. Related to Fig 1.**
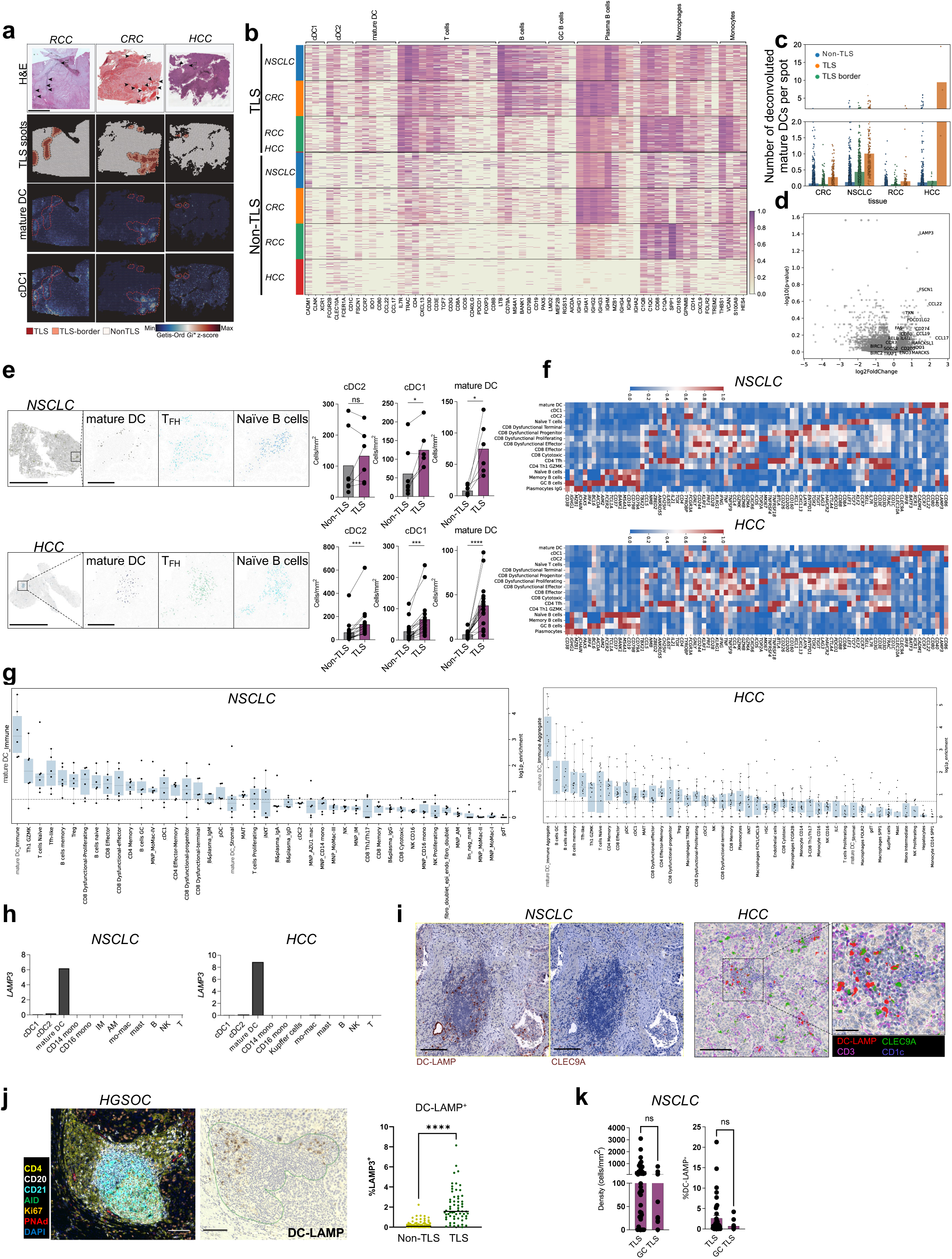
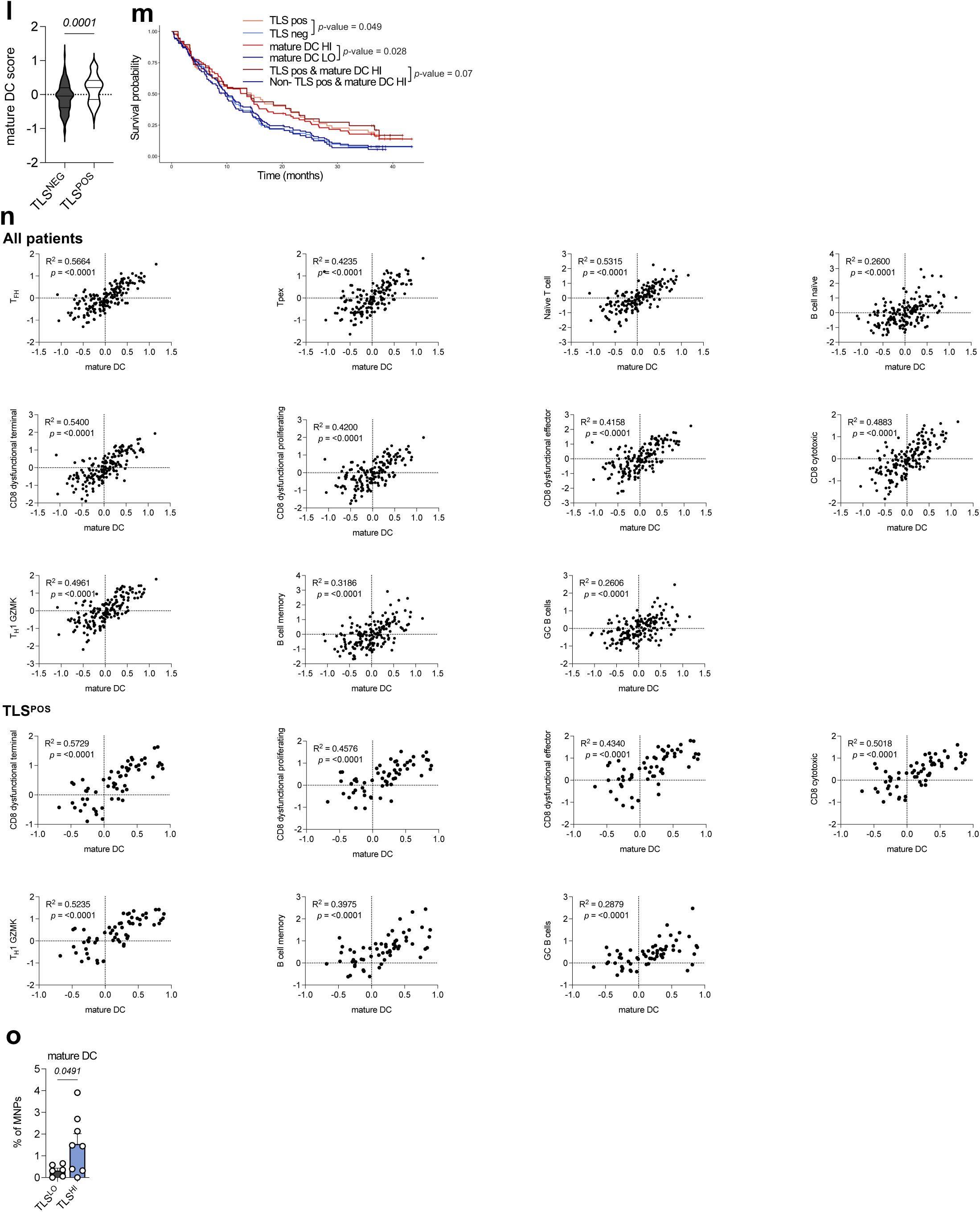
**a-d** Formalin-fixed paraffin-embedded (FFPE) Visium Spatial Transcriptomics analysis of NSCLC (*n* = 2), HCC (*n* = 1), CRC (*n* = 4), and RCC tumors (*n* = 4, data from *Meylan et al., Immunity, 2022*). **a** Representative Visium HCC, CRC and RCC samples showing pathology-annotated TLS, TLS and border spots (two layers of spots located outside the TLS), and hotspots for mature DC and cDC1 gene signatures. Scale bar, 1mm. **b** Truthplot heatmap showing the expression of selected genes representative of DC, T cells, B cells, macrophages, and monocytes in TLS spots compared to non-TLS spots across tumor types. **c** Number of deconvoluted and segmented mature DC per Visium spot in non-TLS (5289, 4670, 1937, and 2103 spots), TLS (3, 128, 187, and 177 spots), and TLS-border spots (14, 177, 770, and 682 spots) across tumor types — respectively HCC, RCC, NSCLC and CRC. **d** Volcano plot comparing TLS mature DC-high Visium spots compared to TLS mature DC-low spots across tumor types. Mature DC gene signature is labeled. **e-g** MERFISH analysis of NSCLC (*n* = 6) and HCC (*n* = 16) tumors. **e** Representative entire TLS area showing deconvoluted and segmented mature DC, T_FH_, and naïve B cells (left). Density of cDC2, cDC1, and mature DC in TLS versus non-TLS areas (right) in NSCLC tumors (top) and HCC tumors (bottom). Scale bars, 5mm (overview) and 200µm (zoom). \**p* < 0.05; \*\*\**p* < 0.001; \*\*\*\**p* < 0.0001 (paired *t*-test). **f** Heatmap showing the expression of selected genes representative of DC, T cells and B cells in NSCLC (top) and HCC (bottom) tumors. **g** Non-curated cell proximity analysis showing enrichment score for immune populations within 30 µm of mature DC in TLS areas in NSCLC (left) and HCC (right). **h** Mean UMI of *LAMP3* expression in immune populations from NSCLC (*n* = 32) and HCC (*n* = 20) tumors analyzed by scRNA-seq. **i** Representative multiplex IHC (MICSSS) images showing mature DC (DC-LAMP^+^) and cDC1 (CLEC9A^+^) in a TLS from an NSCLC tumor (left, representative of 23 patients, scale bar, 200µm) and mature DC, cDC1, cDC2 (CD1c^+^), and T cells (CD3^+^) in a TLS from an HCC tumor (right, representative of 6 patients, scale bars, 200µm (overview) and, 50µm (zoom)). **j** Representative multiplex IF staining of a mature TLS (CD4, CD20, CD21, AID, Ki67, PNAd, DAPI), paired with consecutive DC-LAMP IHC staining of a HGSOC tumor (left). Percentage of mature DC/segmented cells in TLS (*n* = 61 TLS) versus non-TLS (*n* = 116 non-TLS) areas (right). \*\*\*\**p* < 0.0001 (unpaired *t*-test). **k** Density (left) and percentage of mature DC (right) in TLS (*n* = 42 TLS) versus GC^+^ TLS (*n* = 14 GC^+^ TLS) in NSCLC tumors. ns (unpaired *t*-test). **l-n** Bulk transcriptomes paired with TLS annotations in NSCLC (*n* = 168, including 60 TLS^HI^ patients, from Genentech POPLAR dataset). **l** Mature DC score in TLS-positive versus TLS-negative patients. **m** Kaplan-Meier analysis comparing survival probability of TLS-positive versus TLS-negative patients, mature DC-high versus mature DC-low patients, and TLS-positive/mature DC-high patients versus all other patients. **n** Correlations of bulk RNA-seq gene signatures between mature DC and all depicted populations in TLS-positive patients and in all patients. Gene signatures for different immune cell types were generated from prior literature (*Magen et al., 2023*). **o** Percentage of mature DC among mononuclear phagocytes in tumor by scRNA-seq in TLS-high versus TLS-low groups in a neoadjuvant anti-PD-1 trial for HCC patients (*n* = 14 patients).

**Extended Fig 2. Related to Fig 2.**
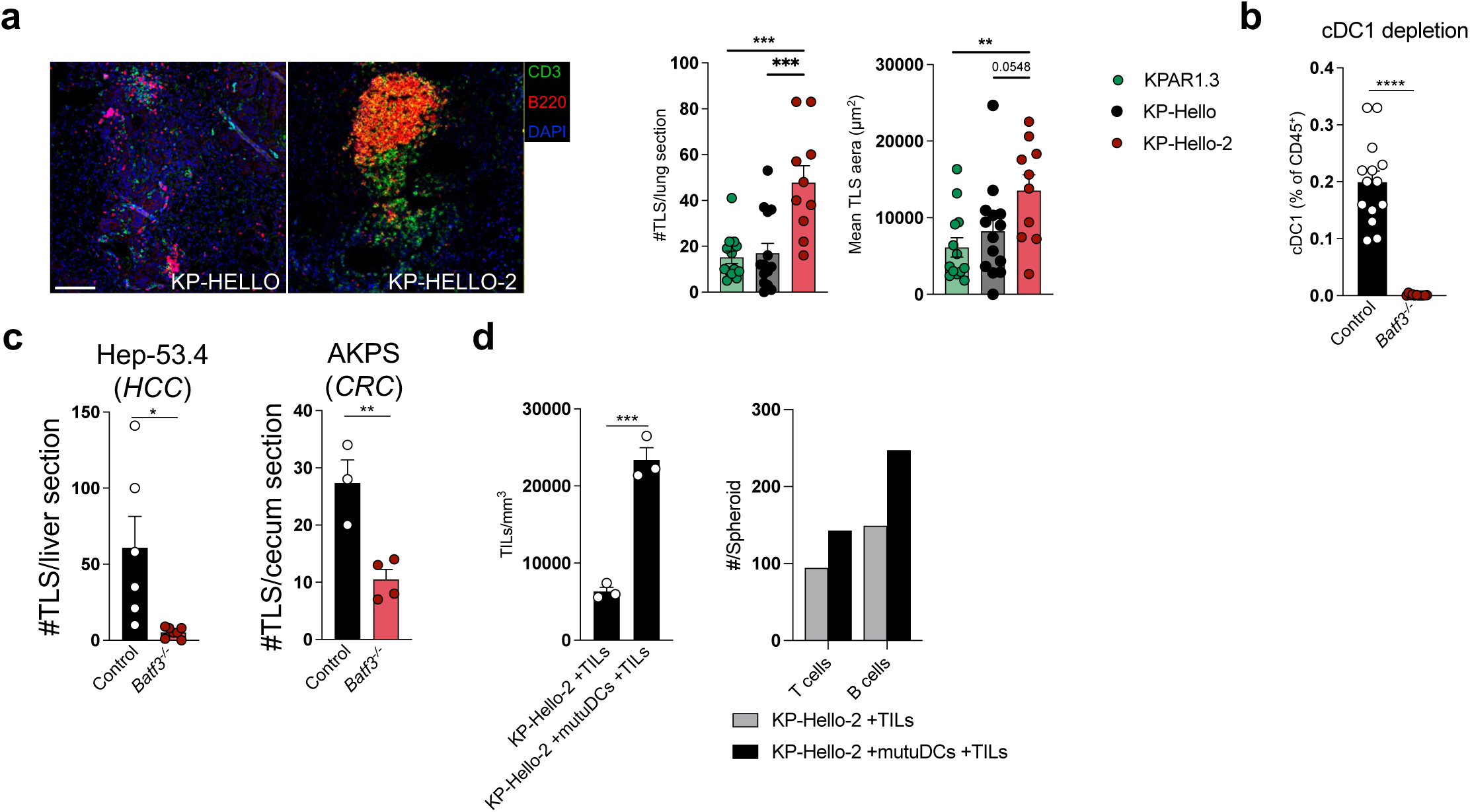
**a** Immunofluorescence analysis of KPAR1.3, KP-HELLO, and KP-HELLO-2 TLS (CD3, B220, DAPI staining), with quantification of TLS number and mean area, at d15 for KP-HELLO and KP-HELLO-2, and at d21 for KPAR1.3 (*n* = 10-14 per group, pooled from two independent experiments). Representative image showing TLS in KP-HELLO and KP-HELLO-2 tumors. Scale bar, 100µm. **b** Intratumor cDC1 depletion efficiency in *Batf3^-/-^*tumor-bearing mice quantified by flow cytometry at d15 (*n* = 11-14 per group, pooled from two independent experiments). **c** TLS number quantification in orthotopic Hep-53.4 HCC tumors (at d15 post-engraftment in the liver, *n* = 6-7 per group) and AKPS CRC tumors (at 6 weeks post-engraftment in the cecum, *n* = 3-4 per group, representative of two independent experiments) in control versus *Batf3^-/-^* mice, based on CD3, B220, and DAPI immunofluorescence staining. **d** KP-HELLO-2 spheroid infiltration and 3D aggregate formation by TILs as measured by BiPhoton microscopy (T and B cells purified from lung tumor-bearing mice at day 7), quantified as the number of TILs per mm³ of spheroid, in the presence or absence of mature mutuDC (cDC1 cell line) previously exposed to apoptotic KP-HELLO-2 cells (*n* = 3 spheroids per condition) (left). Total T and B cell number, pooled across conditions, measured by flow cytometry (*n* = 7 spheroids per condition) (right). \**p* < 0.05; \*\**p* < 0.01; \*\*\**p* < 0.001; \*\*\*\**p* < 0.0001 (unpaired *t*-test).

**Extended Fig 3. Related to Fig 3.**
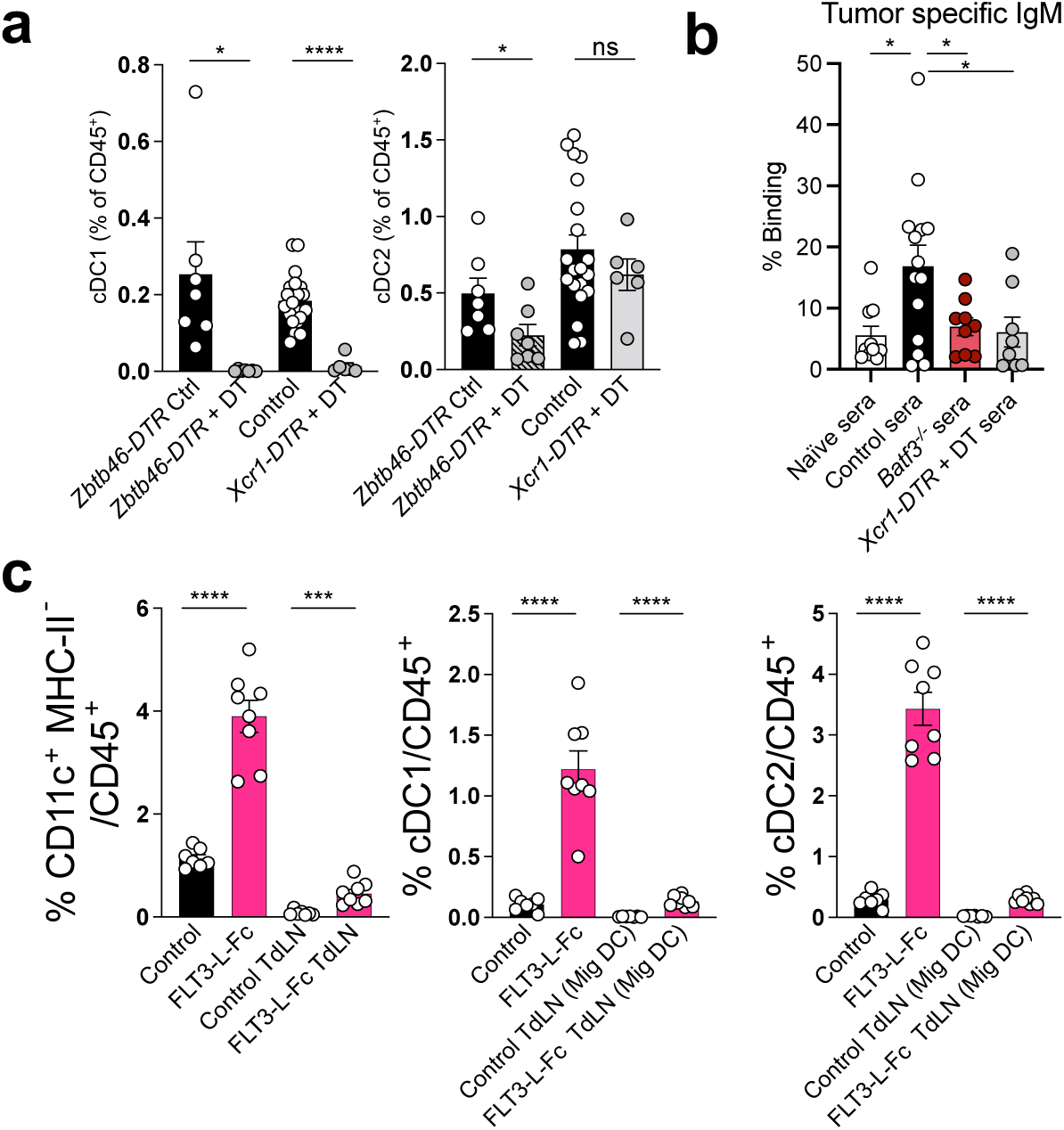
**a** Percentage of intratumor cDC1/CD45^+^ (left) and cDC2/CD45^+^ (right) in *Zbtb46-DTR*, *Xcr1-DTR*, and control mice treated or not with DT, in KP-HELLO-2 tumor-bearing mice, quantified by flow cytometry at d15 (*n* = 6-20 per group, pooled from two independent experiments). **b** Serum IgM binding to tumor cells *ex vivo* from naïve, control, *Batf3-/-*, and *Xcr1-DTR* mice treated with DT (*n* = 8-14 per group, pooled from two independent experiments). Sera were collected at d15. **c** Percentage of Lineage^−^ MoMac^−^ CD11c^+^ MHC-II^−^/CD45^+^ (left), cDC1/CD45^+^ (middle) and cDC2/CD45^+^ (right) in tumor-bearing lungs and tdLN (migratory DC), in WT mice treated or not with FLT3-L-Fc at day 8, quantified by flow cytometry at d15 (*n* = 7-8 per group, representative of two independent experiments). \**p* < 0.05; \*\*\**p* < 0.001; \*\*\*\**p* < 0.0001 (unpaired *t*-test).

**Extended Fig 4. Related to Fig. 4.**
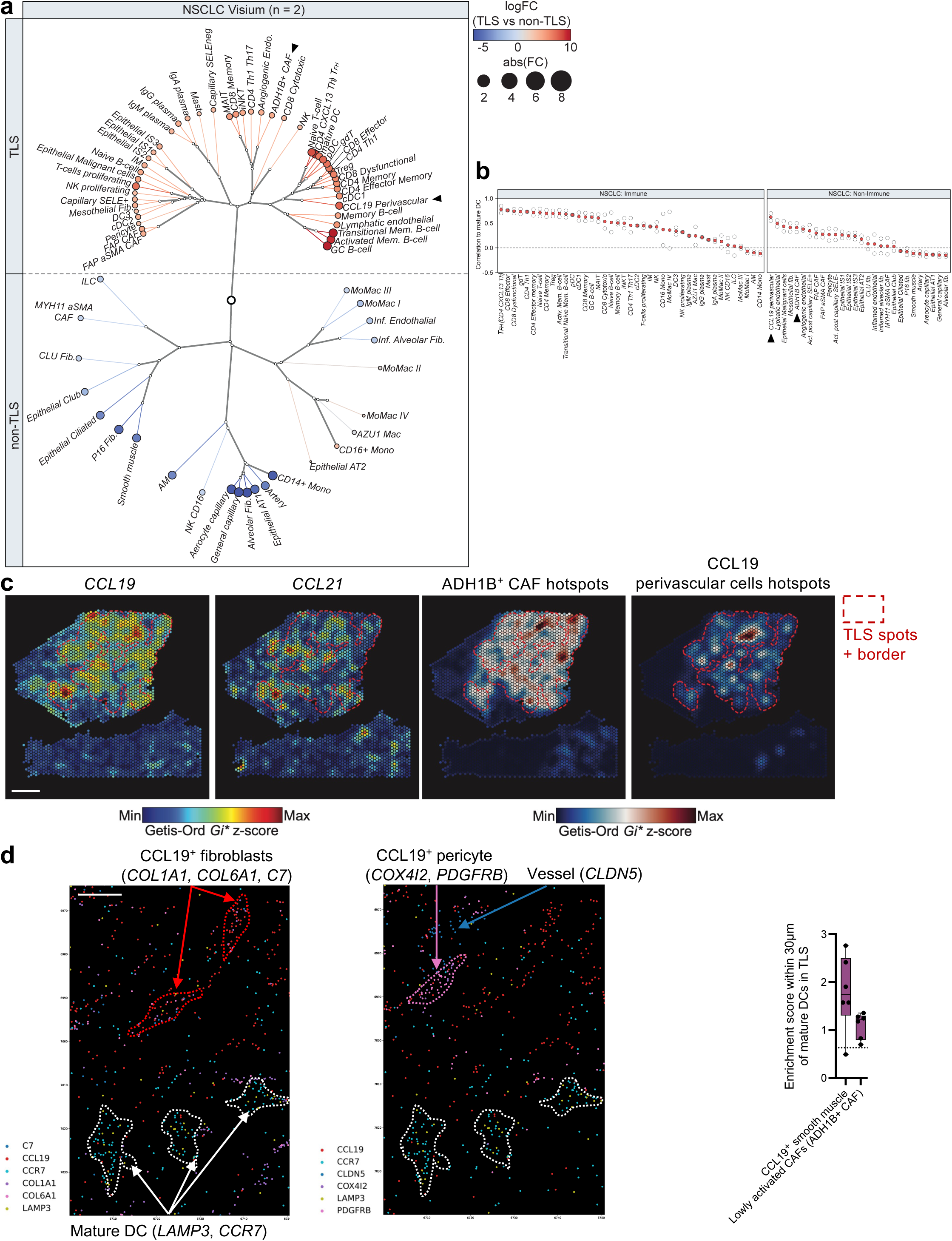
**a-c** Formalin-fixed paraffin-embedded (FFPE) Visium Spatial Transcriptomics analysis of NSCLC (*n* = 2) **a** Hierarchical clustering tree (Daylight Tree) representing the co-enrichment of cell types, displaying log fold change (logFC) and absolute fold change in TLS spots versus non-TLS spots across NSCLC immune and non-immune cell types, deconvoluted from scRNA-seq references and segmented. **b** Spatial correlation with mature DC among immune and non-immune cell types. Arrows indicate ADH1B^+^ CAFs and CCL19^+^ perivascular cells. **c** Representative Visium NSCLC sample showing pathology-annotated TLS and border spots, along with hotspots for *CCL19* and *CCL21* gene expression and ADH1B^+^ CAF and CCL19^+^ perivascular cell gene signatures. Scale bar, 1mm. **d** MERFISH Spatial Transcriptomics analysis of NSCLC tumors. Representative TLS area illustrating the proximity between mature DC and *CCL19* positive stromal cells (*n* = 1) (left). Scale bar, 10µm. Cell proximity analysis showing the enrichment score for ADH1B^+^ CAFs and CCL19^+^ perivascular cells within 30 µm of mature DC in TLS areas (*n* = 6) (right).

**Extended Fig 5. Related to Fig. 4.**
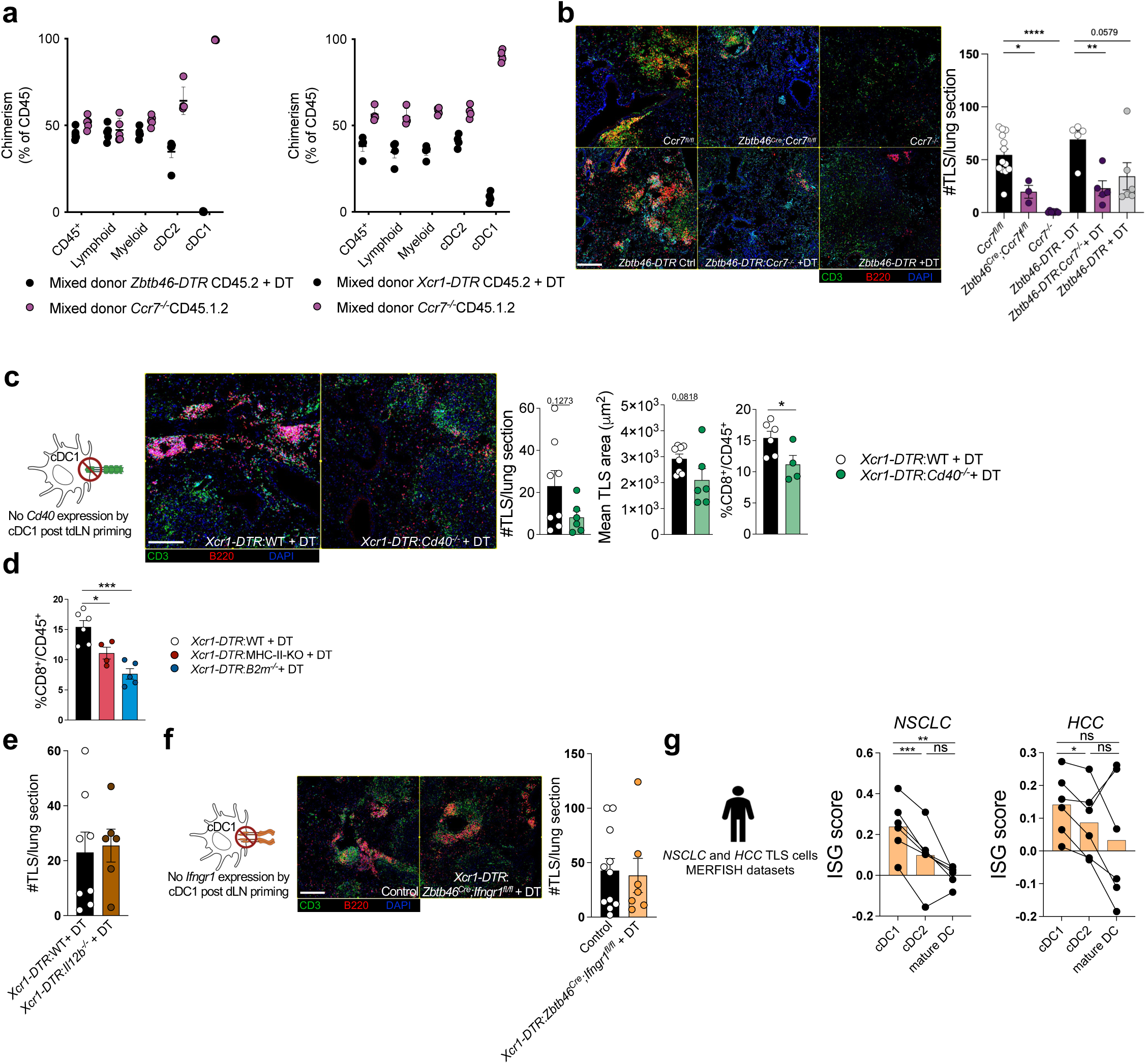
**a** Chimerism of donor bone marrow cells (CD45.2^+^ or CD45.1.2^+^) expressed as a percentage of total CD45^+^ cells in the indicated populations in DT-treated mixed bone marrow chimeras *Zbtb46-DTR* CD45.2:*Ccr7^-/-^* CD45.1.2 mice (left, *n* = 4 per group) and *Xcr1-DTR* CD45.2:*Ccr7^-/-^* CD45.1.2 mice (right, *n* = 5 per group, representative of two independent experiments) at d15 post-tumor engraftment. **b-c, e, f** Immunofluorescence analysis of TLS (stained for CD3, B220, and DAPI) and quantification of TLS number at d15 post-tumor engraftment. **b** Representative images (left) and quantification in *Ccr7^fl/fl^*, *Zbtb46^Cre^;Ccr7^fl/fl^*, *Ccr7^-/-^* mice, in *Zbtb46-DTR* bone marrow chimera mice treated or not with DT, and *Zbtb46-DTR* CD45.2:*Ccr7^-/-^*CD45.1.2 mixed bone marrow chimera mice treated with DT (*n* = 3-13 per group, representative of two independent experiments). Scale bar, 200µm. **c** Quantification of TLS number and mean area in DT-treated *Xcr1- DTR:*WT versus *Xcr1-DTR:Cd40^-/-^* mixed bone marrow chimera mice (n = 6-8 per group, scale bar, 200µm) and intratumor quantification by flow cytometry at d15 of CD8^+^ T cells among immune cells. **d** Intratumor quantification by flow cytometry at d15 of CD8^+^ T cells among immune cells in DT-treated *Xcr1-DTR*/WT, *Xcr1- DTR*:MHC-II-KO and *Xcr1-DTR:B2m^-/-^* mixed bone marrow chimera mice (*n* = 4-6 per group, representative of two independent experiments). **b-d** \**p* < 0.05; \*\**p* < 0.01; \*\*\**p* < 0.005; \*\*\*\**p* < 0.001 (unpaired *t*-test). **e** TLS number in DT-treated *Xcr1-DTR:*WT versus *Xcr1-DTR:Il12b^-/-^* mixed bone marrow chimera mice (*n* = 6- 8 per group). **f** Representative image and TLS number in control bone marrow chimera versus DT-treated *Xcr1-DTR:Zbtb46^Cre^;Ifngr1^fl/fl^*mixed bone marrow chimera mice (*n* = 7-11 per group). Scale bar, 200µm. **g** MERFISH spatial transcriptomics analysis of human NSCLC (*n* = 6, left) and HCC (*n* = 6, right) tumors showing normalized ISG gene module scores for cDC1, cDC2, and mature DC within TLS. \**p* < 0.05; \*\**p* < 0.01; \*\*\**p* < 0.005; (paired *t*-test).

Please, see Excel files for Supplemental Tables:

- Supplemental-Table-1-Patient_Clinical_Metadata
- Supplemental-Table-2-MERFISH_Gene_Panels
- Supplemental-Table-3-Antibodies
- Supplemental-Table-4-Gene_List_Bulk_Seq
- Supplemental-Table-5-Flow_Cytometry_Gating_Strategy

