## Supplemental-Table-1-Patient_Clinical_Metadata for "Dendritic cells type 1 control the formation, maintenance, and function of tertiary lymphoid structures in cancer"

| ID | Gender | Age | Pack-Year | Histologic Type | T stage | N stage | M stage | TLS high | Recurrence | TNM stage |
|---|---|---|---|---|---|---|---|---|---|---|
| 338 | F | 74 | 13 | Lung Adenocarcinoma | 1b | 0 | 0 | Yes | NA | NA |
| 370 | F | 69 | 0 | Lung Adenocarcinoma | 2a | 1 | x | Yes | NA | NA |
| 377 | M | 77 | 25 | Lung Squamous-Cell Carcinoma | 3 | 0 | x | Yes | NA | NA |
| 403 | M | 67 | 10 | Lung Adenocarcinoma | 3 | 0 | x | Yes | NA | NA |
| 408 | M | 79 | 45 | Lung Adenocarcinoma | 2b | 0 | x | Yes | NA | NA |
| 410 | F | 76 | 35 | Lung Adenocarcinoma | 1b | 0 | x | Yes | NA | NA |
| 514 | F | 66 | 30 | Lung Adenocarcinoma | 3 | 0 | x | Yes | NA | NA |
| 522 | F | 82 | 30 | Lung Squamous-Cell Carcinoma | 2a | 0 | x | Yes | NA | NA |
| 558 | F | 57 | 20 | Lung Adenocarcinoma | 4 | 2 | x | Yes | NA | NA |
| 570 | M | 76 | 0 | Lung Adenocarcinoma | 2b | 0 | x | Yes | NA | NA |
| 571 | M | 64 | 0 | Lung Adenocarcinoma | 1c | 0 | x | Yes | NA | NA |
| 578 | F | 71 | 40 | Lung Adenocarcinoma | 1a | 0 | x | Yes | NA | NA |
| 581 | M | 68 | 0 | Lung Adenocarcinoma | 2a | 0 | x | Yes | NA | NA |
| 706 | M | 74 | 125 | Lung Squamous-Cell Carcinoma | 2b | 0 | x | Yes | NA | NA |
| 371 | M | 79 | 28 | Lung Adenocarcinoma | 1b | x | x | No | NA | NA |
| 378 | F | 64 | 13 | Lung Adenocarcinoma | 1b | 0 | x | No | NA | NA |
| 393 | F | 62 | 10 | Lung Adenocarcinoma | 1b | 0 | x | No | NA | NA |
| 458 | F | 90 | 0 | Lung Adenocarcinoma | 1c | 0 | x | No | NA | NA |
| 460 | M | 65 | 0 | Lung Adenocarcinoma | 1b | 0 | x | No | NA | NA |
| 464 | F | 70 | 20 | Lung Adenocarcinoma | 1b | 0 | x | No | NA | NA |
| 532 | M | 73 | 1.7 | Lung Adenocarcinoma | 1c | 0 | x | No | NA | NA |
| 564 | F | 67 | 48 | Lung Adenocarcinoma | 1b | 0 | x | No | NA | NA |
| 569 | F | 61 | 4 | Lung Adenocarcinoma | 1a | 0 | x | No | NA | NA |
| 572 | M | 64 | 10 | Lung Adenocarcinoma | 1b | x | x | No | NA | NA |
| 584 | M | 72 | 40 | Lung Squamous-Cell Carcinoma | 2a | 1 | x | No | NA | NA |
| 593 | F | 75 | 47 | Lung Adenocarcinoma | 1b | 0 | x | No | NA | NA |
| 596 | M | 65 | 30 | Lung Adenocarcinoma | 1b | x | x | No | NA | NA |
| 626 | F | 56 | 0 | Lung Adenocarcinoma | 1c | 0 | x | No | NA | NA |
| 630 | M | 66 | 35 | Lung Adenocarcinoma | 1b | 0 | x | No | NA | NA |
| 695 | F | 73 | 15 | Lung Adenocarcinoma | 2b | 0 | x | No | NA | NA |
| 714 | F | 76 | 75 | Lung Squamous-Cell Carcinoma | 2a | 1 | NA | No | NA | NA |
| 725 | F | 64 | >0 | Lung Adenocarcinoma | 1b | 0 | x | No | NA | NA |
| BIC18 | F | 54 | NA | Colon Adenocarcinoma | 2 | 2a | x | NA | NA | NA |
| BIC21 | F | 77 | NA | Colon Adenocarcinoma | 3 | 0 | x | NA | NA | NA |
| BIC31 | M | 54 | NA | Colon Adenocarcinoma | 3 | 2b | x | NA | NA | NA |
| 1003 | M | 46 | NA | Hepatocellular Carcinoma | NA | NA | NA | Yes | Yes | II |
| 1005 | F | 79 | NA | Hepatocellular Carcinoma | NA | NA | NA | No | Yes | II |
| 1006 | M | 48 | NA | Hepatocellular Carcinoma | NA | NA | NA | Yes | Yes | IIIA |
| 1014 | F | 83 | NA | Hepatocellular Carcinoma | NA | NA | NA | Yes | No | NA |
| 1017 | M | 72 | NA | Hepatocellular Carcinoma | NA | NA | NA | Yes | No | NA |
| 1023 | M | 64 | NA | Hepatocellular Carcinoma | NA | NA | NA | Yes | No | NA |
| 1024 | M | 46 | NA | Hepatocellular Carcinoma | NA | NA | NA | No | No | IB |
| 1025 | M | 67 | NA | Hepatocellular Carcinoma | NA | NA | NA | No | Yes | II |
| 1029 | M | 68 | NA | Hepatocellular Carcinoma | NA | NA | NA | Yes | No | II |
| 1030 | M | 66 | NA | Hepatocellular Carcinoma | NA | NA | NA | No | No | IA |
| 1031 | M | 73 | NA | Hepatocellular Carcinoma | NA | NA | NA | No | NA | NA |
| 1032 | M | 55 | NA | Hepatocellular Carcinoma | NA | NA | NA | No | Yes | IIIA |
| 1037 | M | 68 | NA | Hepatocellular Carcinoma | NA | NA | NA | Yes | No | NA |
| 1039 | F | 71 | NA | Hepatocellular Carcinoma | NA | NA | NA | No | Yes | II |
| 1041 | M | 73 | NA | Hepatocellular Carcinoma | NA | NA | NA | Yes | NA | IB |
