## Supplemental-Table-2-MERFISH_Gene_Panels for "Dendritic cells type 1 control the formation, maintenance, and function of tertiary lymphoid structures in cancer"

| MERSCOPE Gene Panel 1 | MERSCOPE Gene Panel 2 |
| --- | --- |
| **Gene** | **Gene** |
| ACTA2 | A4GALT |
| ACTG2 | ACTA2 |
| ADAM12 | ACTG2 |
| ADAM28 | ADAM12 |
| ADGRE5 | ADH1B |
| ADH1B | AICDA |
| CCL21 | AIM2 |
| AIM2 | ANKRD55 |
| ANKRD55 | AREG |
| AREG | ATF3 |
| ASPN | ATP1A2 |
| ATP1A2 | BAAT |
| BAAT | BANK1 |
| BANK1 | BATF |
| BATF3 | BATF3 |
| BGN | BCL2 |
| BIRC3 | BCL2L1 |
| BRD2 | BCL6 |
| BTG2 | BGN |
| BTG3 | BIRC3 |
| BTLA | BST2 |
| C1QA | BTLA |
| C1QB | C1QA |
| C1QC | C1QB |
| C4BPA | C1QC |
| CADM1 | C4BPA |
| CALM2 | CADM1 |
| CAPG | CAPG |
| CASQ2 | CASP8 |
| CCDC50 | CASQ2 |
| CCDC80 | CCL17 |
| CCL18 | CCL18 |
| CCL2 | CCL19 |
| CCL22 | CCL2 |
| CCND1 | CCL20 |
| CCND2 | CCL22 |
| CCNL1 | CCL4 |
| CCR4 | CCL5 |
| CCR6 | CCL7 |
| CCR7 | CCL8 |
| CCR8 | CCND1 |
| CCRL2 | CCND2 |
| CD14 | CCR2 |
| CD160 | CCR3 |
| CD163 | CCR4 |
| CD19 | CCR5 |
| CD1C | CCR6 |
| CD1D | CCR7 |
| CD1E | CCR8 |

CD2
CD200
CD247
CD27
CD28
CD300E
CD37
CD38
CD3E
CD4
CD40
CD40LG
CD44
CD53
CD59
CD5L
CD6
CD69
CD7
CD74
CD79A
CD79B
CD80
CD82
CD83
CD86
CD8A
CD8B
CD9
CD99
CDKN1A
CDKN1C
CENPF
CETP
CFLAR
CFP
CH25H
CHMP1B
CLDN5
CLEC10A
CLEC4C
CLEC9A
CMC1
CNN1
COL1A1
COL1A2
COL3A1
COL4A1
COL4A2
COL6A1
COMP
CPVL

CCRL2
CD14
CD160
CD163
CD164
CD19
CD1C
CD1D
CD1E
CD2
CD200
CD207
CD209
CD22
CD226
CD244
CD248
CD27
CD274
CD276
CD28
CD34
CD36
CD37
CD38
CD3D
CD3E
CD3G
CD4
CD40
CD40LG
CD44
CD5
CD5L
CD6
CD68
CD69
CD7
CD70
CD74
CD79A
CD79B
CD80
CD83
CD86
CD8A
CD8B
CD9
CDH1
CDH2
CDH3
CDH4

CREM
CRTAM
CSF1R
CTHRC1
CTLA4
CX3CR1
CXCL10
CXCL11
CXCL12
CXCL13
CXCL3
CXCL8
CXCL9
CXCR3
CXCR4
CXCR5
CXCR6
CYCS
CYP27A1
CYTIP
DCN
DES
DIO2
DNASE1L3
DPT
DUSP4
DUSP5
DYNLL1
EIF4A3
ELF1
ENTPD1
EOMES
EREG
ERICH1
ESM1
EZR
F13A1
FAM3C
TENT5C
FAP
FBP1
FCER1A
FCGRT
FCN1
FHL5
FLT3
FN1
FOLR2
FOS
FOSB
FOXP3
FPR2

CDH5
CDK2
CDK4
CDK6
CEACAM8
CEBPB
CENPF
CETP
CH25H
CLEC10A
CLEC14A
CLEC4C
CLEC7A
CLEC9A
CNN1
COL11A1
COL1A1
COL1A2
COL3A1
COL4A1
COL5A1
CR1
CR2
CSF1
CSF1R
CSF2
CSF2RA
CSF3
CSF3R
CTGF
CTLA4
CTNNB1
CTSH
CX3CL1
CX3CR1
CXCL1
CXCL10
CXCL12
CXCL13
CXCL16
CXCL2
CXCL3
CXCL5
CXCL8
CXCL9
CXCR1
CXCR2
CXCR3
CXCR4
CXCR5
CXCR6
DNMT3A

| | |
|---|---|
| FSCN1 | DPP4 |
| FUCA1 | DUSP1 |
| GATA3 | EGF |
| GBP1 | EGFR |
| GBP5 | ELN |
| GJB2 | ENTPD1 |
| GLT8D2 | EOMES |
| GLUL | EPCAM |
| GNG4 | EREG |
| GPC3 | ESAM |
| GPNMB | F13A1 |
| GPR171 | FABP2 |
| GPR183 | FABP3 |
| GREM1 | FABP4 |
| GRN | FAP |
| GSN | FAS |
| GUCY1A2 | FASLG |
| GZMK | FBP1 |
| H3F3B | FCER2 |
| HAVCR2 | FCGR2A |
| HERPUD1 | FCGR3A |
| HES1 | FCGRT |
| HMGB2 | FCN1 |
| HMOX1 | FCRL6 |
| JPT1 | FLT1 |
| HOPX | FLT3 |
| HSD17B6 | FLT3LG |
| ICA1 | FLT4 |
| ICAM1 | FOLR2 |
| ICOS | FOS |
| IDO1 | FOXP3 |
| IDS | FSCN1 |
| IER2 | GATA3 |
| IFI44L | GLYCAM1 |
| IFNGR1 | GNLY |
| IFT57 | GPC3 |
| IGFBP4 | GPNMB |
| IGHM | GPR183 |
| IL10 | GPR65 |
| IL12A | GSN |
| IL12B | GZMA |
| IL15 | GZMB |
| IL18BP | GZMH |
| IL18R1 | GZMK |
| IL18RAP | HAVCR2 |
| IL1B | HES1 |
| IL1R2 | HIF1A |
| IL1RN | HLA-A |
| IL21 | HLA-B |
| IL21R | HLA-C |
| IL2RA | HLA-DMB |
| IL2RB | HLA-DQA1 |

IL33
IL3RA
IL6ST
IL7R
INHBA
IRF4
IRF7
IRF8
ISG20
ITGAE
ITGB1
ITGB2
ITIH3
ITM2A
JCHAIN
JUN
JUNB
KCNA5
KIF2A
KIT
KLF2
KLF3
KLF6
KLRB1
KLRG1
LAG3
LAMP3
LDB3
LEF1
LGALS3
LGMN
LILRA4
LILRB5
LIPA
LITAF
LMNA
LRRC15
LSP1
LUM
LY9
LYST
LYVE1
LYZ
MAF
MAFB
MARCKSL1
MARCO
MASP2
MCL1
MC1R
MGAT4A
MGP

HLA-DQA2
HLA-DQB1
HLA-DQB2
HLA-DRA
HLA-DRB1
HLA-DRB5
HMGB2
HSD17B6
ICAM1
ICAM2
ICAM3
ICOS
ICOSLG
IDO1
IDO2
IFI6
IFIT1
IFITM1
IFITM10
IFITM2
IFITM5
IFNAR1
IFNAR2
IFNB1
IFNG
IFNGR1
IFNGR2
IGHA2
IGHG1
IGHMI
IL12A
IL12B
IL12RB2
IL13
IL15
IL17A
IL18
IL18BP
IL18R1
IL18RAP
IL1A
IL1B
IL1R1
IL1R2
IL1RN
IL2
IL21
IL21R
IL22
IL23A
IL2RA
IL2RB

| | |
|---|---|
| MKI67 | IL33 |
| MMP2 | IL3RA |
| MRC1 | IL4 |
| MS4A1 | IL4I1 |
| MXRA5 | IL5RA |
| MYADM | IL6 |
| MYH11 | IL6R |
| NCR3 | IL6ST |
| NEK6 | IL7R |
| NFKB1 | IRF1 |
| NFKBIA | IRF3 |
| NFKBIZ | IRF4 |
| NLRP3 | IRF5 |
| NR4A1 | IRF7 |
| NR4A2 | IRF8 |
| NTRK3 | ISG15 |
| NUB1 | ISG20 |
| OLFML3 | ITGA1 |
| OSM | ITGA4 |
| P4HA3 | ITGA5 |
| PAMR1 | ITGAE |
| PAX5 | ITGAL |
| DRAIC | ITGAM |
| PDCD1 | ITGAV |
| PDE4B | ITGAX |
| PDE4D | ITGB1 |
| PDGFRA | ITGB2 |
| PDGFRB | ITIH3 |
| PDGFRL | JAK1 |
| PDK4 | JCHAIN |
| PECAM1 | JUN |
| PLAC8 | KIR2DL4 |
| PLAUR | KIR3DL2 |
| PLEK | KIT |
| PLN | KLF2 |
| PLP2 | KLF6 |
| PLTP | KLRB1 |
| PLVAP | KLRC1 |
| PNRC1 | KLRD1 |
| PON1 | KLRF1 |
| POSTN | KLRG1 |
| POU2F2 | KLRK1 |
| PPP1R15A | KRAS |
| PRDM1 | KRT10 |
| PRELP | KRT18 |
| PRF1 | KRT19 |
| PROX1 | KRT20 |
| PSAP | KRT7 |
| PTGS2 | L10 |
| PTPN1 | LAG3 |
| RAB9A | LAIR1 |
| RASGEF1B | LAMP3 |

| | |
|---|---|
| RBPJ | LAYN |
| RGS5 | LEF1 |
| RHOB | LGALS3 |
| RHOC | LGALS9 |
| RNASE1 | LGMN |
| RORA | LILRA4 |
| S100B | LILRB5 |
| S1PR1 | LIPA |
| S1PR5 | LMNA |
| SCIMP | LOX |
| SDC1 | LTA |
| SDCBP | LTB |
| SELE | LY6E |
| SELL | LYVE1 |
| SELENOP | LYZ |
| SERPIND1 | MADCAM1 |
| SESN3 | MAF |
| SFRP2 | MAFB |
| SH2D2A | MARCO |
| SIRPG | MGP |
| SLC2A3 | MICA |
| SLC40A1 | MICB |
| SMAP2 | MKI67 |
| SMIM14 | MME |
| SNAP47 | MMP1 |
| SNX3 | MMP11 |
| SNX9 | MMP12 |
| SOCS3 | MMP2 |
| SOX4 | MMP7 |
| SPON1 | MMP9 |
| SPP1 | MMRN1 |
| SRSF5 | MMRN2 |
| SRSF7 | MPO |
| STAB1 | MRC1 |
| STAT3 | MS4A1 |
| STK17A | MT2A |
| STK17B | MTOR |
| STMN1 | MUC1 |
| SUB1 | MUC2 |
| SUGCT | MUC4 |
| SVEP1 | MX1 |
| TACSTD2 | MYC |
| TAGLN | MYH11 |
| TAGLN2 | MZB1 |
| TCF21 | NCAM1 |
| TCF4 | NCR1 |
| TCF7 | NCR3 |
| TCL1A | NFKB1 |
| TFF3 | NFKB2 |
| THBS1 | NFKBIA |
| THBS2 | NKG7 |
| THEMIS2 | NLRP3 |

TIGIT
TIMD4
CEMIP2
TNF
TNFAIP2
TNFAIP3
TNFRSF13B
TNFRSF8
TNFSF13B
TNFSF8
TOP2A
TOX
TOX2
TRAC
TREM1
TREM2
TRGC1
TSC22D3
TSPAN13
TSPYL2
TUBB
TXNIP
UBC
UGT2B4
VCAM1
VCAN
VPS37B
VWF
WARS
WDFY4
CCN4
CCN5
XBP1
XCR1
YPEL5
ZBTB16
ZC3HAV1
ZFP36L1
ZNF331

NMB
NOS2
NOS3
NOTCH1
NR4A1
NR4A2
NRAS
NTAN1
OAS1
OSM
PAX5
PDCD1
PDCD1LG2
PDGFA
PDGFB
PDGFC
PDGFRA
PDGFRB
PDPN
PECAM1
PIK3CA
PIK3CG
PLAC8
PLAUR
PLTP
PLVAP
PRDM1
PRF1
PROX1
PSMB8
PSMB9
PTPRC
RAG1
RGMB
RHOB
RNASE1
RTP4
S100A10
S100A12
S100A4
S100A8
S100A9
SCIMP
SDC1
SELL
SELP
SELP-1
SELPLG
SEMA3A
SERPIND1
SERPINE1
SIRPG

SLBP
SLC40A1
SLC4A10
SOCS1
SOCS3
SOX2
SOX4
SOX9
SPP1
STAB1
STAP1
STAT1
STAT3
STAT4
STAT5A
STAT6
SVEP1
TAP1
TAP2
TBX21
TCF21
TCF4
TCF7
TCL1A
TET2
TGFB1
TGFB2
TGFB3
TGFBI
TGFBR1
TGFBR2
TGFBR3
THBS1
THBS2
TIGIT
TIMD4
TLR1
TLR2
TLR4
TLR9
TMEM173
TNF
TNFAIP2
TNFRSF13B
TNFRSF13C
TNFRSF14
TNFRSF17
TNFRSF18
TNFRSF4
TNFRSF9
TNFSF10
TNFSF11

TNFSF13B
TNFSF14
TNFSF18
TNFSF4
TNFSF9
TOP2A
TOX
TOX2
TP53
TP63
TRAC
TRBC1
TRDC
TREM2
TRGC1
TUBB
TYROBP
VCAM1
VCAN
VEGFA
VEGFB
VEGFC
VWF
WNT3
WNT3A
WNT5A
WT1
WWTR1
XBP1
XCL1
XCR1
YAP1
ZAP70
ZBED2
ZEB1
