## Supplemental-Table-3-Antibodies for "Dendritic cells type 1 control the formation, maintenance, and function of tertiary lymphoid structures in cancer"

Supplemental Table 2: Antibodies

*Anti-mouse antibodies (note: all antibodies were used at 1:200 unless otherwise stated; human antibodies are specified whereapplicable)

| Target | Clone | Fluorochrome | Catalog # | Company | Dilution | Use | Reactivity |
|---|---|---|---|---|---|---|---|
| B220 | RA3-6B2 | e450 | 48-0452-82 | eBioscience | | Flow Cytometry | |
| B220 | RA3-6B2 | BV711 | 103255 | Biolegend | | Flow Cytometry | |
| BCL6 | IG191E/A8 | PE | 648303 | Biolegend | 1/100 | Flow Cytometry | |
| CCR7 | 4B12 | PE | 120105 | Biolegend | 1/100 | Flow Cytometry | |
| CCR7 | 4B12 | PE-Cy7 | 120124 | Biolegend | 1/100 | Flow Cytometry | |
| CD119 | 2E2 | PE | 12-1191-82 | eBioscience | 1/100 | Flow Cytometry | |
| CD11b | M1/70 | PerCP-Cy5.5 | 101230 | Biolegend | | Flow Cytometry | |
| CD11b | M1/70 | e450 | 48-0112-82 | eBioscience | | Flow Cytometry | |
| CD11b | M1/70 | PE-Cy7 | 25-0112-82 | eBioscience | | Flow Cytometry | |
| CD11b | M1/70 | APC-R700 | 564985 | BD | | Flow Cytometry | |
| CD11c | N418 | APC-e780 | 47-0114-82 | eBioscience | | Flow Cytometry | |
| CD11c | N418 | BV785 | 117336 | Biolegend | | Flow Cytometry | |
| CD19 | eBio1D3 | e450 | 48-0193-82 | eBioscience | | Flow Cytometry | |
| CD19 | eBio1D3 | PerCP-Cy5.5 | 45-0193-82 | eBioscience | | Flow Cytometry | |
| CD19 | 6D5 | Alexa700 | 115528 | Biolegend | | Flow Cytometry | |
| CD19 | 1D3 | BV650 | 563235 | BD | | Flow Cytometry | |
| CD25 | PC61 | PerCP-Cy5.5 | 102028 | Biolegend | | Flow Cytometry | |
| CD39 | 24DMS1 | PE-Cy7 | 25-0391-82 | eBioscience | | Flow Cytometry | |
| CD3ε | 17A2 | BV510 | 100233 | Biolegend | | Flow Cytometry | |
| CD3ε | eBio500A2 | FITC | 11-0033-82 | eBioscience | | Flow Cytometry | |
| CD3ε | 145-2C11 | PE | 553064 | BD | | Flow Cytometry | |
| CD3ε | 145-2C11 | PerCP-Cy5.5 | 45-0031-82 | eBioscience | | Flow Cytometry | |
| CD3ε | 17A2 | APC-e780 | 47-0032-82 | eBioscience | | Flow Cytometry | |
| CD3ε | 17A2 | PerCP-eFluor710 | 46-0032-82 | eBioscience | | Flow Cytometry | |
| CD3ε | 500A2 | AF700 | 557984 | BD | | Flow Cytometry | |
| CD3ε | 145-2C11 | Biotin | 13-0031-85 | eBioscience | | Flow Cytometry | |
| CD4 | GK1.5 | PE-Cy5 | 100409 | Biolegend | | Flow Cytometry | |
| CD4 | RM4-5 | BV650 | 100545 | Biolegend | | Flow Cytometry | |
| CD40 | 1C10 | APC | 17-0401-82 | eBioscience | | Flow Cytometry | |
| CD44 | RM4-5 | BV605 | 100548 | Biolegend | | Flow Cytometry | |
| CD44 | IM7 | PE-Cy7 | 25-0441-82 | eBioscience | | Flow Cytometry | |
| CD44 | IM7 | A700 | 103026 | Biolegend | | Flow Cytometry | |
| CD45 | 30-F11 | BV650 | 103151 | Biolegend | | Flow Cytometry | |
| CD45 | 30-F11 | BV510 | 103138 | Biolegend | | Flow Cytometry | |
| CD45 | 30-F11 | BV605 | 103139 | Biolegend | | Flow Cytometry | |
| CD45.1 | A20 | FITC | 4323293 | BD | | Flow Cytometry/Multiplex IF | |
| CD45.1 | A20 | APC | 17-0453-82 | eBioscience | | Flow Cytometry | |
| CD45.1 | A20 | BV510 | 110741 | Biolegend | | Flow Cytometry | |
| CD45.2 | 104 | FITC | 11-0454-82 | eBioscience | | Flow Cytometry | |
| CD45.2 | 104 | APC-e780 | 47-0454-82 | eBioscience | | Flow Cytometry | |
| CD62L | MEL-14 | FITC | 104406 | Biolegend | | Flow Cytometry | |
| CD64 | X54-5/7.1 | APC | 17-0641-82 | eBioscience | | Flow Cytometry | |
| CD64 | X54-5/7.1 | Biotin | 139318 | Biolegend | | Flow Cytometry | |
| CD69 | H1.2F3 | BV711 | 104537 | Biolegend | | Flow Cytometry | |
| CD8α | 53-6.7 | BV510 | 100751 | Biolegend | | Flow Cytometry | |
| CD8α | 53-6.7 | FITC | 553031 | BD | | Flow Cytometry | |
| CD8α | 53-6.7 | APC | 100711 | Biolegend | | Flow Cytometry | |
| CD8α | 53-6.7 | APC-e780 | 100714 | Biolegend | | Flow Cytometry | |
| CD8α | 53-6.7 | PerCP-Cy5.5 | 100732 | Biolegend | | Flow Cytometry | |
| CD80 | 16-10A1 | PE-Cy5 | 15-0801-81 | eBioscience | | Flow Cytometry | |
| CD86 | GL1 | FITC | 105006 | Biolegend | | Flow Cytometry | |
| CD86 | GL1 | Biotin | 13-0862-82 | eBioscience | | Flow Cytometry | |
| CD86 | GL-1 | Pacific Blue | 105022 | Biolegend | | Flow Cytometry | |
| CXCR5 | L138D7 | BV650 | 145517 | Biolegend | 1/100 | Flow Cytometry | |
| F4/80 | BM8 | AF488 | 123120 | Biolegend | | Flow Cytometry | |
| F4/80 | BM8 | Biotin | 123105 | Biolegend | | Flow Cytometry | |
| F4/80 | BM8 | BV605 | 123133 | Biolegend | | Flow Cytometry | |
| FAS | SA367H8 | FITC | 152605 | Biolegend | | Flow Cytometry | |
| FoxP3 | FJK-16s | PE | 12-5773-82 | eBioscience | 1/100 | Flow Cytometry | |
| FoxP3 | MF-14 | Alexa700 | 126422 | Biolegend | 1/100 | Flow Cytometry | |
| GL7 | GL7 | Alexa647 | 561529 | BD | | Flow Cytometry/Multiplex IF | |
| GL7 | GL7 | FITC | 562080 | BD | | Flow Cytometry | |
| Goat anti-mouse IgG | Poly4053 | PE-Cy7 | 405315 | Biolegend | | Flow Cytometry | |
| Gr-1 (Ly6G/Ly6C) | RB6-8C5 | Biotin | 13-5931-81 | eBioscience | | Flow Cytometry | |
| H-2Kb | AF6-88.5.5.3 | ef450 | 48-5958-82 | eBioscience | 1/200 | Flow Cytometry | |
| ICOS | C398.4A | A488 | 313514 | Biolegend | | Flow Cytometry | |
| IFNg | XMG1.2 | Alexa700 | 557998 | BD | 1/100 | Flow Cytometry | |
| IgD | 11-26c-2a | BV711 | 405731 | Biolegend | | Flow Cytometry | |
| IgM | II/41 | PerCP-eFluor710 | 46-5790-82 | eBioscience | | Flow Cytometry | |
| IgM | RMM-1 | BV711 | 406539 | , Biolegend | | Flow Cytometry | |
| Ki-67 | SolA15 | FITC | 11-5698-82 | eBioscience | 1/100 | Flow Cytometry | |
| Ly6C | HK1.4 | BV605 | 128035 | Biolegend | | Flow Cytometry | |
| Ly6G | 1A8 | AF700 | 127622 | Biolegend | | Flow Cytometry | |
| MerTK | 2B10C42 | PE | 151506 | Biolegend | 1/100 | Flow Cytometry | |
| MHC-II (I-A/I-E) | M5/114.15.2 | AF700 | 56-5321-82 | eBioscience | 1/400 | Flow Cytometry | |
| MHC-II (I-A/I-E) | M5/114.15.2 | BV711 | 107643 | Biolegend | 1/400 | Flow Cytometry | |
| NK1.1 | PK136 | PE-Cy7 | 108714 | Biolegend | | Flow Cytometry | |
| NK1.1 | PK136 | Biotin | 13-5941-85 | eBioscience | | Flow Cytometry | |
| NK1.1 | PK136 | Alexa700 | 560515 | BD | | Flow Cytometry | |
| PD-1 | J43 | PE-Cy7 | 25-9985-82 | eBioscience | | Flow Cytometry | |
| PD-1 | 29F.1A12 | BV785 | 135225 | Biolegend | | Flow Cytometry | |

| | | | | | | | |
|---|---|---|---|---|---|---|---|
| Streptavidin | | PerCP-Cy5.5 | 45-4317-82 | eBioscience | 1/500 | Flow Cytometry | |
| Streptavidin | | BV650 | 405232 | Biolegend | 1/500 | Flow Cytometry | |
| T-bet | 4B10 | Pacific Blue | 644807 | Biolegend | 1/100 | Flow Cytometry | |
| TCF1 | C63D9 | AF488 | 6444S | Cell Signaling Technology | 1/100 | Flow Cytometry | |
| TCF1 | C63D9 | Pacific Blue | 9066S | Cell Signaling Technology | 1/100 | Flow Cytometry | |
| Tim-3 | RMT3-23 | BV605 | 119721 | Biolegend | | Flow Cytometry | |
| TNFa | MP6-XT22 | APC | 506307 | Biolegend | 1/100 | Flow Cytometry | |
| XCR1 | ZET | BV650 | 148220 | Biolegend | | Flow Cytometry | |
| XCR1 | ZET | BV785 | 148225 | Biolegend | | Flow Cytometry | |
| XCR1 | ZET | APC-Cy7 | 148223 | Biolegend | | Flow Cytometry | |
| CD11c | D1V9Y | | 97585S | Cell Signaling Technology | 1/100 | Multiplex IHC | |
| DsRed/cross react to mCherry | Polyclonal Rabbit IgG | | 632496 | Takara Bio | | Multiplex IHC | |
| B220 | RA3-6B2 | | 14-0452-82 | eBioscience | | Multiplex IF | |
| CD3 | Polyclonal Rabbit IgG | | 17617-1-AP | Proteintech | | Multiplex IF | |
| CD3 | Polyclonal Rabbit IgG | | PA5-32318 | Invitrogen | | Multiplex IF | |
| CD8a | 4SM15 | | 14-0808-82 | eBioscience | 1/100 | Multiplex IF | |
| PD-1 | Polyclonal Goat IgG | | AF1021 | R&D Systems | 1/100 | Multiplex IF | |
| TCF1/7 | C63D9 | | 2203T | Cell Signaling Technology | 1/100 | Multiplex IF | |
| CD4 | EPR19514 | | ab183685 | Abcam | 1/1000 | Multiplex IF | |
| FITC | AB_2339038 | Alexa488 | 200-542-037 | Jackson ImmunoResearch | | Multiplex IF | |
| Ki67 | SP6 | Opal 520 | MA5-14520 | ThermoFisher | 1/100 | Multiplex IF | |
| CD21 | SP186 | Opal 620 | ab227662 | Abcam | 1/50 | Multiplex IF | |
| AID | mAID-2 | Opal 690 | 14-5959-82 | ThermoFisher | 1/800 | Multiplex IF | |
| Goat Anti-Rabbit | Polyclonal Goat IgG | Alexa488 | R37116 | Invitrogen | 1/500 | Multiplex IF | |
| Goat Anti-Rat | Polyclonal Goat IgG | Alexa647 | A-21247 | Invitrogen | 1/500 | Multiplex IF | |
| Donkey Anti-Rat | Polyclonal Donkey IgG | Alexa647 | A-78947 | Invitrogen | 1/500 | Multiplex IF | |
| Donkey Anti-Rabbit | Polyclonal Donkey IgG | Alexa594 | R37119 | Invitrogen | 1/500 | Multiplex IF | |
| Donkey Anti-Goat | Polyclonal Donkey IgG | Alexa488 | A-11055 | Invitrogen | 1/500 | Multiplex IF | |
| Donkey Anti-Goat | Polyclonal Donkey IgG | Alexa647 | ab150131 | Abcam | 1/500 | Multiplex IF | |
| Donkey Anti-Rat | Polyclonal Donkey IgG | Alexa594 | A-21209 | Invitrogen | 1/500 | Multiplex IF | |
| CCR7 | 4B12 | Alexa488 | 120110 | Biolegend | 1/100 | CyCIF | |
| CD86 | E5W6H | Alexa647 | 19589 | Cell Signaling Technology | 1/100 | CyCIF | |
| CD4 | 4SM95 | eFluor 570 | 41-9766-82 | eBioscience | | CyCIF | |
| CD8a | D4V8L | Alexa647 | 83012BC | Cell Signaling Technology | | CyCIF | |
| Bcl-6 | D-8 | Alexa647 | sc-7388 AF647 | Santa Cruz | | CyCIF | |
| TCF1/7 | C63D9 | Alexa488 | 6444S | Cell Signaling Technology | | CyCIF | |
| CD45R | RA3-6B2 | eFluor 570 | 41-0452-80 | eBioscience | | CyCIF | |
| Ki-67 | D3B5 | Alexa647 | 12075 | Cell Signaling Technology | | CyCIF | |
| CD11c | D1V9Y | Alexa 555 | 64675BC | Cell Signaling Technology | | CyCIF | |
| MHCII | 78593BC | Alexa647 | 78593BC | Cell Signaling Technology | | CyCIF | |
| DC-LAMP | 1010E1.01 | | DDX0191P-100 | Novus biologicals | 1/80 | Multiplex IHC | Human |
| CLEC9A | EPR22324 | | ab223188 | Abcam | 1/1000 | Multiplex IHC | Human |
| CD3 | 2GV6 | | 790-4341 | Ventana | RTU | Multiplex IHC | Human |
| CD1c | OTI2F4 | | ab156708 | Abcam | 1/150 | Multiplex IHC | Human |
| ADH1B | OTI3F2 | | MA5-25547 | ThermoFisher | 1/500 | Multiplex IHC | Human |
| CCL19 | Polyclonal Goat IgG | | AF361 | R&D systems | 1/150 | Multiplex IHC | Human |
| CD20 | L26 | | M0755 | Dako | 1/250 | Multiplex IHC | Human |
| MYH11 | JA03-35 | | NBP2-66967 | Novus biologicals | 1/300 | Multiplex IHC | Human |
| CD4 | EP204 | | API3209AA | Biocare | | Multiplex IF | Human |
| CD20 | L26 | | NC-L-CD20-L26 | Leica | | Multiplex IF | Human |
| CD21 | 2G9 | | NCL-L-CD21-2G9 | Leica | | Multiplex IF | Human |
| AID | EPR23436-45 | | ab269454 | Abcam | | Multiplex IF | Human |
| Ki67 | SP6 | | RM9106S | Epredia (Fisher) | | Multiplex IF | Human |
| PNAd | MECA-79 | | MABF2050 | Sigma | | Multiplex IF | Human |
