## Supplemental-Table-4-Gene_List_Bulk_Seq for "Dendritic cells type 1 control the formation, maintenance, and function of tertiary lymphoid structures in cancer"

Supplemental Table 3: Gene List for Bulk Seq Analysis

| | | | | | | | | | | | | | | | | | | | | | | | | | | | |
|---|---|---|---|---|---|---|---|---|---|---|---|---|---|---|---|---|---|---|---|---|---|---|---|---|---|---|---|
| mature DC | *CCR7* | *CCL17* | *CCL19* | *CCL22* | *IL12B* | *MARCKS* | *MARCKSL1* | *CD80* | *TRAF1* | *RELB* | *BIRC2* | *BIRC3* | *ENO3* | *LAMP3* | *FSCN1* | *CD274* | *PDCD1LG2* | *CD200* | *FAS* | *SOCS2* | *IL4I1* | *IDO1* | *TXN* | | | | |
| T naïve | *TRAC* | *CD3D* | *CD3E* | *CD3G* | *IL7R* | *SELL* | *CCR7* | *KLF2* | *TCF7* | *LEF1* | | | | | | | | | | | | | | | | | |
| CD8 Dysfunctional Terminal | *TRAC* | *CD3D* | *CD3E* | *CD3G* | *CD8A* | *CD8B* | *PDCD1* | *CTLA4* | *HAVCR2* | *LAG3* | *TIGIT* | *TOX2* | *ENTPD1* | *LAYN* | | | | | | | | | | | | | |
| CD8 Dysfunctional Progenitor (Tpex) | *TRAC* | *CD3D* | *CD3E* | *CD3G* | *CD8A* | *CD8B* | *PDCD1* | *CXCL13* | *DUSP4* | *XCL1* | *XCL2* | *CD160* | *CD200* | *CD226* | *BTLA* | *GNG4* | *TNFRSF18* | *TNFRSF4* | | | | | | | | | |
| CD8 Dysfunctional Proliferating | *TRAC* | *CD3D* | *CD3E* | *CD3G* | *CD8A* | *CD8B* | *PDCD1* | *MKI67* | *TOP2A* | | | | | | | | | | | | | | | | | | |
| CD8 Dysfunctional Effector | *TRAC* | *CD3D* | *CD3E* | *CD3G* | *CD8A* | *CD8B* | *PDCD1* | *ICOS* | *CXCR6* | *GZMA* | *GZMB* | *GZMK* | | | | | | | | | | | | | | | |
| CD8 Effector | *TRAC* | *CD3D* | *CD3E* | *CD3G* | *CD8A* | *CD8B* | *CCL3L3* | *CCL4* | *CCL4L2* | *TNFSF9* | *TNF* | *IFNG* | *KLRG1* | *CD28* | | | | | | | | | | | | | |
| CD8 Cytotoxic | *TRAC* | *CD3D* | *CD3E* | *CD3G* | *CD8A* | *CD8B* | *PRF1* | *CMC1* | *KLRF1* | *CD244* | *GNLY* | *FCGR3A* | *TYROBP* | | | | | | | | | | | | | | |
| Tfh | *TRAC* | *CD3D* | *CD3E* | *CD3G* | *CD4* | *PDCD1* | *CTLA4* | *TIGIT* | *TOX* | *TOX2* | *ICA1* | *CXCL13* | *DUSP4* | *BTLA* | *CD200* | *IL21* | *IL6ST* | *CH25H* | *ANKRD55* | *ZBED2* | *GNG4* | *NMB* | *LHFP* | *EBI3* | *PCAT29* | *ICOS* | *CD28* |
| Th1-GZMK | *TRAC* | *CD3D* | *CD3E* | *CD3G* | *CD4* | *ICOS* | *CD28* | *IL10* | *GZMK* | *GZMA* | *CCL4L2* | *CCL5* | *TNFSF9* | *TNF* | *CXCR6* | *IFNG* | *TBX21* | *CCL4* | | | | | | | | | |
| Naïve B cells | *CD79A* | *CD79B* | *CD19* | *MS4A1* | *BANK1* | *CD37* | *TCL1A* | *FCER2* | *IGHD* | *IGHM* | | | | | | | | | | | | | | | | | |
| Memory B cells | *CD79A* | *CD79B* | *CD19* | *MS4A1* | *BANK1* | *CD37* | *AIM2* | | | | | | | | | | | | | | | | | | | | |
| GC B cells | *CD79A* | *CD79B* | *CD19* | *MS4A1* | *BANK1* | *LMO2* | *MEF2B* | *RGS13* | *AICDA* | *BCL6* | | | | | | | | | | | | | | | | | |
