## Supplemental-Table-5-Flow_Cytometry_Gating_Strategy for "Dendritic cells type 1 control the formation, maintenance, and function of tertiary lymphoid structures in cancer"

| | cDC1 | cDC2 | Mig cDC1 (tdLN) | Mig cDC2 (tdLN) | CD11c$^{+}$ MHC-II$^{-}$ | Tfh | Th1 | CD8$^{+}$ T cells | CD8$^{+}$ PD-1+ T cells | Tpex | CD8$^{+}$ PD-1+ CD39+ T cells | GC B cells |
|---|---|---|---|---|---|---|---|---|---|---|---|---|
| **Gated on** | Lin- Mono- AM- Mac- cDC2- | Lin- Mono- AM- Mac- cDC1- | Lin- Mono- Mac- cDC2- | Lin- Mono- Mac- cDC1- | Lin- Mono- AM- Mac- | B cell- CD8- | B cell- CD8- Treg- | B cell- CD4- | B cell- CD4- | B cell- CD4- | B cell- CD4- | T cell- Naïve B cell- |
| **LiveDead** | - | - | - | - | - | - | - | - | - | - | - | - |
| **CD45** | + | + | + | + | + | + | + | + | + | + | + | + |
| **CD19** | - (Lin) | - (Lin) | - (Lin) | - (Lin) | - (Lin) | - | - | - | - | - | - | + |
| **CD3ε** | - (Lin) | - (Lin) | - (Lin) | - (Lin) | - (Lin) | + | + | + | + | + | + | - |
| **NK1.1** | - (Lin) | - (Lin) | - (Lin) | - (Lin) | - (Lin) | | | | | | | |
| **Ly-6G** | - (Lin) | - (Lin) | - (Lin) | - (Lin) | - (Lin) | | | | | | | |
| **Ly-6C** | - | - | - | - | | | | | | | | |
| **CD64** | - | - | - | - | - | | | | | | | |
| **F4/80** | - | - | - | - | - | | | | | | | |
| **CD11c** | + | + | + | + | + | | | | | | | |
| **MHC-II** | + | + | hi | hi | - | | | | | | | |
| **CD11b** | - | + | - | + | | | | | | | | |
| **XCR1** | + | - | + | - | | | | | | | | |
| **CD4** | | | | | | + | + | - | - | - | - | |
| **CD8α** | | | | | | - | - | + | + | + | + | |
| **PD-1** | | | | | | + | | | + | + | + | |
| **CXCR5** | | | | | | + | | | | | | |
| **FoxP3** | | | | | | | - | | | | | |
| **IFNg** | | | | | | | or + | | | | | |
| **T-bet*** | | | | | | | or + | | | | | |
| **CD39** | | | | | | | | | | - | + | |
| **TCF1** | | | | | | | | | | + | - | |
| **IgD** | | | | | | | | | | | | - |
| **igM** | | | | | | | | | | | | - |
| **GL7** | | | | | | | | | | | | + |

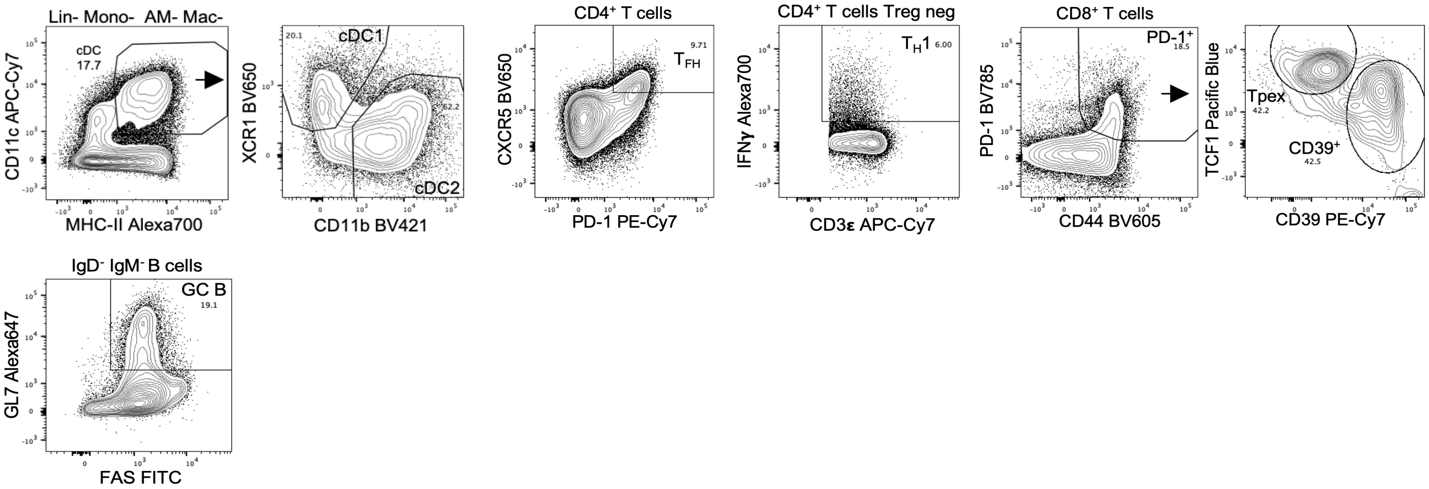
